# RBM-5 modulates U2AF large subunit-dependent alternative splicing in *C. elegans*

**DOI:** 10.1101/355388

**Authors:** Chuanman Zhou, Xiaoyang Gao, Surong Hu, Wenjing Gan, Xu Jing, Ma Yong-Chao, Ma Long

## Abstract

A key step in pre-mRNA splicing is the recognition of 3’ splicing sites by the U2AF large and small subunits, a process regulated by numerous *trans*-acting splicing factors. How these *trans*-acting factors interact with U2AF *in vivo* is unclear. From a screen for suppressors of the temperature-sensitive (ts) lethality of the *C. elegans* U2AF large subunit gene *uaf-1(n4588)* mutants, we identified mutations in the <u>R</u>NA <u>b</u>inding <u>m</u>otif gene *rbm-5*, a homolog of the tumor suppressor *RBM5*. *rbm-5* mutations can suppress *uaf-1(n4588)* ts-lethality by loss of function and neuronal expression of *rbm-5* was sufficient to rescue the suppression. Transcriptome analyses indicate that *uaf-1(n4588)* affected the expression of numerous genes and *rbm-5* mutations can partially reverse the abnormal gene expression to levels similar to that of wild type. Though *rbm-5* mutations did not obviously affect alternative splicing per se, they can suppress or enhance, in a gene-specific manner, the altered splicing of genes in *uaf-1(n4588)* mutants. Specifically, the recognition of a weak 3’ splice site was more susceptible to the effect of *rbm-5*. Our findings provide novel *in vivo* evidence that RBM-5 can modulate UAF-1-dependent RNA splicing and suggest that RBM5 might interact with U2AF large subunit to affect tumor formation.

**Author summary:** RNA splicing is a critical regulatory step for eukaryotic gene expression and has been involved in the pathogenesis of multiple diseases. How RNA splicing factors interact *in vivo* to affect the splicing and expression of genes is unclear. In studying the temperature-sensitive lethal phenotypes of a mutation affecting the splicing factor U2AF large subunit gene *uaf-1* in the nematode *Caenorhabditis elegans*, we isolated suppressive mutations in the *rbm-5* gene, a homolog of the human tumor suppressor gene *RBM5*. *rbm-5* is broadly expressed in neurons to enhance the lethality of the *uaf-1* mutants. We found that the uaf-1 mutation causes aberrant expression of genes in numerous biological pathways, a large portion of which can be corrected by *rbm-5* mutations. The abnormal splicing of multiple genes caused by the *uaf-1* mutation is either corrected or enhanced by *rbm-5* mutations in a gene-specific manner. We propose that RBM-5 interacts with UAF-1 to affect RNA splicing and the tumor suppressor function of RBM5 might involve U2AF-dependent RNA splicing.

## Introduction

Pre-mRNA splicing (RNA splicing) is a highly regulated process that generates mature mRNAs in eukaryotic cells by removing intervening non-coding introns from pre-mRNA transcripts and simultaneously joins coding exons [1, 2]. RNA splicing is carried out in a coordinated two-step trans-esterification reaction that is catalyzed by the spliceosome, a complex of small nuclear ribonucleoproteins (snRNPs) [1–3], and facilitated by hundreds of splicing factors [4]. Using different 5’ or 3’ splice sites and by including or skipping introns and exons from a single form of pre-mRNA transcript, alternative splicing can generate multiple distinct mRNA isoforms, which increases the proteome size of an organism and is considered a driving force for the biological complexity of metazoans [1, 5, 6]. Defects in RNA splicing are the causes of numerous human diseases [7, 8].

RNA splicing are affected by numerous *trans* splice factors, *cis* splice elements, transcription, chromatin structures and histone modifications [1, 4, 9]. Hundreds of protein factors related to RNA splicing have been identified [10–13]. However, it is unclear how these factors interact to affect the splicing of various genes.

RNA recognition motif (RRM) is an abundant protein domain that can bind both RNAs and proteins [14]. The <u>R</u>NA-<u>b</u>inding <u>m</u>otif protein 5 gene (*RBM5/LUCA-15/H37*) encodes a protein containing two RRMs, two zinc fingers, one arginine-serine-rich domain (RS) and one glycine patch (G-patch) [15]. *RBM5* and its paralog *RBM6* are located in the human chromosome 3p21.3 region and were identified as candidate tumor suppressors of lung cancers and other solid tumors [16–22]. *RBM10*, another paralog of *RBM5*, is also frequently mutated in lung adenocarcinoma samples [23]. In mice and human, *RBM5* can reduce lung cancer progression [24, 25].

RBM5 can increase the expression of proapoptotic protein BAX and reduce that of anti-apoptotic proteins including BCL-2 and BCL-XL [26–28]. It could also affect the expression of other apoptosis and cell cycle genes [29]. RBM5 might regulate the splicing of apoptosis-and cancer-related genes by affecting the recognition of 3’ splice sites (3’ SS) [30–33]. The *in vivo* mechanism underlying the function of RBM5 is unclear.

*C. elegans* is a model organism for studying a variety of biological processes [34]. We previously identified mutations affecting the *C. elegans* orthologs of the U2AF large subunit (UAF-1 in *C. elegans*, U2AF65 or U2AF2 in mammals), splicing factor one (SFA-1) [35] and the splicing factor microfibrillar associated protein 1 (MFAP-1) [36] by screening for essential gene suppressors of a *unc-93(gf)* rubberband phenotype [37]. Studies of these mutants provided new insights into the in vivo regulation of alternative splicing [35, 36, 38] and suggest a previously unknown interaction of *uaf-1* with the *C. elegans* spinal muscular atrophy gene *smn-1* in affecting locomotion and lifespan [39]. The *uaf-1(n4588)* mutation causes temperature-sensitive (ts) lethality [35]. A screen for suppressors of the ts-lethal phenotype identified intragenic suppressors of *uaf-1* and potentially extragenic suppressors [35].

In this study, we describe the genetic and molecular characterization of two extragenic suppressors of *uaf-1(n4588).* We found that they affect the *C. elegans rbm-5*. We also analyzed how *rbm-5* and *uaf-1* interact to affect gene expression and alternative splicing.

## Results

### The *uaf-1(n4588)* mutation causes embryonic lethality and sterility at high temperatures

We previously isolated the *uaf-1(n4588)* missense mutation from a screen for essential gene suppressors of the “rubberband” Unc phenotype caused by the *unc-93(e1500)* gain-of-function mutation [35]. *uaf-1(n4588)* mutants were inviable at 25 °C and grew like wild type at 15 °C [35]. At 20 °C, *uaf-1(n4588)* mutants grew slightly slower and have partial protruding vulva (Pvl) and/or sterility (Ste) phenotypes (Fig. S1). We also found that *uaf-1(n4588)* mutants were inviable at 22.5 °C (this study).

To examine how temperatures affect the embryonic development of *uaf-1(n4588)* mutants, eggs laid at 15 °C were shifted to 20 °C, 22.5 °C or 25 °C. With these treatments, most eggs would hatch in 24 hrs (Fig. 1A). We next shifted L4 *uaf-1(n4588)* mutants grown at 15 °C to higher temperatures, allowing them to grow into adults and lay eggs for 24 hrs. Most eggs laid at 20 °C and 22.5 °C would hatch, while only ~10% eggs laid at 25 °C did so (Fig. 1B).

**Figure 1.**
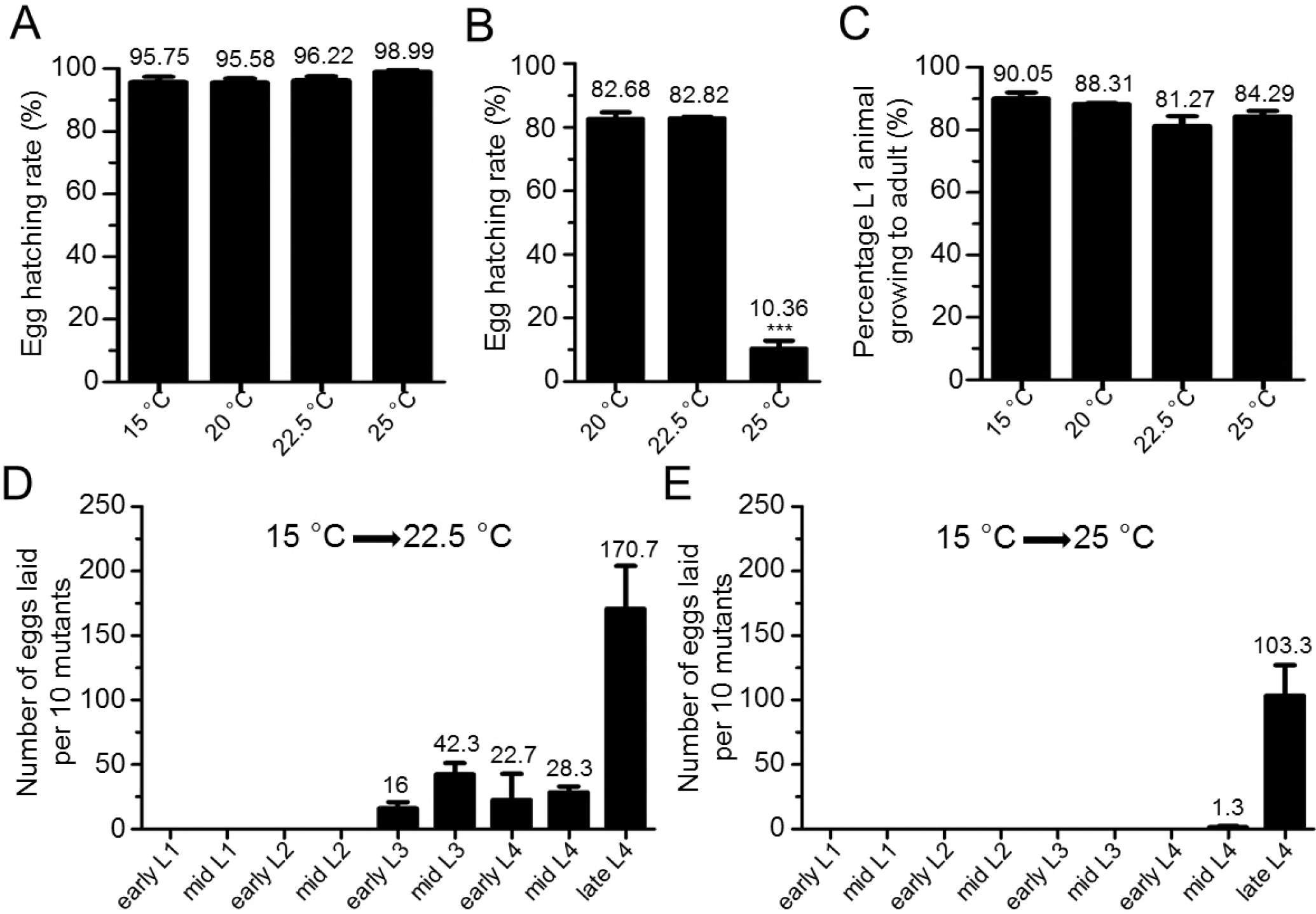
*uaf-1(n4588)* mutants exhibited embryonic lethality and sterility at high temperatures. (A) *uaf-1(n4588)* mutant eggs laid at 15 °C were shifted to higher temperatures and the hatching rates were quantified. (B) L4 *uaf-1(n4588)* animals grown at 15 °C where shifted to higher temperatures, allowed to lay eggs for 24 hrs, and the egg-hatching rates were quantified. (C) L1 *uaf-1(n4588)* animals grown at 15 °C were shifted to higher temperatures and the percentages of animals that grew to adults were quantified. (D, E) *uaf-1(n4588)* mutants of various developmental stages grown at 15 °C were shifted to 22.5 °C or 25 °C, respectively, and allowed to develop to the mid L4 stage. 10 L4 animals were picked to new plates and allowed to grow and lay eggs for 24 hrs. The numbers of eggs were counted. Results were from 3 to 5 biological replicates. For A, B and C, 90 to 150 eggs or L1 animals were analyzed in each biological replicate. The value of each column is on top. Statistics: paired two-tailed Student’s t-test. *: p<0.05; **: p<0.01; ***: p<0.001.

To understand how high temperatures affect the larval development of *uaf-1(n4588)* mutants, we shifted L1 larva grown at 15 °C to higher temperatures, at which most larvae could grow into adults (Fig. 1C). However, adults growing up at 22.5 °C and 25 °C were sterile.

To determine whether high temperatures affect specific developmental stages in causing sterility, we grew *uaf-1(n4588)* mutants to various larval stages at 15 °C and shifted them to 22.5 °C or 25 °C, allowing the larvae to grow into adults and lay eggs. When shifted to 22.5 °C, L2 or younger larvae would become sterile adults (Fig. 1D), while at 25 °C mid-L4 or younger larvae would become so (Fig. 1E). Together these results suggest that *uaf-1(n4588)* might cause lethality by affecting both embryonic development and fertility at high temperatures.

### Extragenic mutations can suppress the sterility and embryonic lethality of *uaf-1(n4588)* mutants at high temperatures

A previous screen for suppressors of *uaf-1(n4588)* ts-lethality at 25 °C identified four intragenic suppressors of *uaf-1* [35]. From the screen we also isolated seven potential suppressors that might be caused by extragenic mutations [35]. When reexamined, two isolates (*n5130*, *n5131*) exhibited extremely slow growth and severe sterility and therefore were not further analyzed.

We examined the five healthier extragenic suppressors (*n5119*, *n5121*, *n5122*, *n5128* and *n5132*) in more details. At all temperatures, these mutations can strongly suppress the egg-laying defect of *uaf-1(n4588*) mutants (Fig. 2A, 2B, 2C). At 20 °C, *uaf-1(n4588)* did not cause an apparent egg-hatching defect and the hatching rates were similar between *uaf-1(n4588)* single and *uaf-1(n4588); sup* double mutants (Fig. 2A’). At 22.5 °C, *uaf-1(n4588)* caused a slightly defective egg-hatching rate, which was suppressed by *n5119*, *n5121*, *n5122* and *n5132* (Fig. 2B’). At 25 °C, *uaf-1(n4588)* mutants had a severe egg-hatching defect, which was strongly suppressed by all five mutations (Fig. 2C’). In addition, the Pvl and Ste phenotypes of *uaf-1(n4588)* mutants were also strongly suppressed by these mutations at 20 °C (Fig. S1).

At 22.5 °C, *uaf-1(n4588)* single mutants would become sterile, while *uaf-1(n4588); sup* double mutants were fertile and could propagate. At 25 °C, *uaf-1(n4588); sup* double mutants could not propagate but appeared to healthier than *uaf-1(n4588)* single mutants. Therefore, we isolated these suppressors probably as escapers in the original screen, which was performed at 25 °C [35].

**Figure 2.**
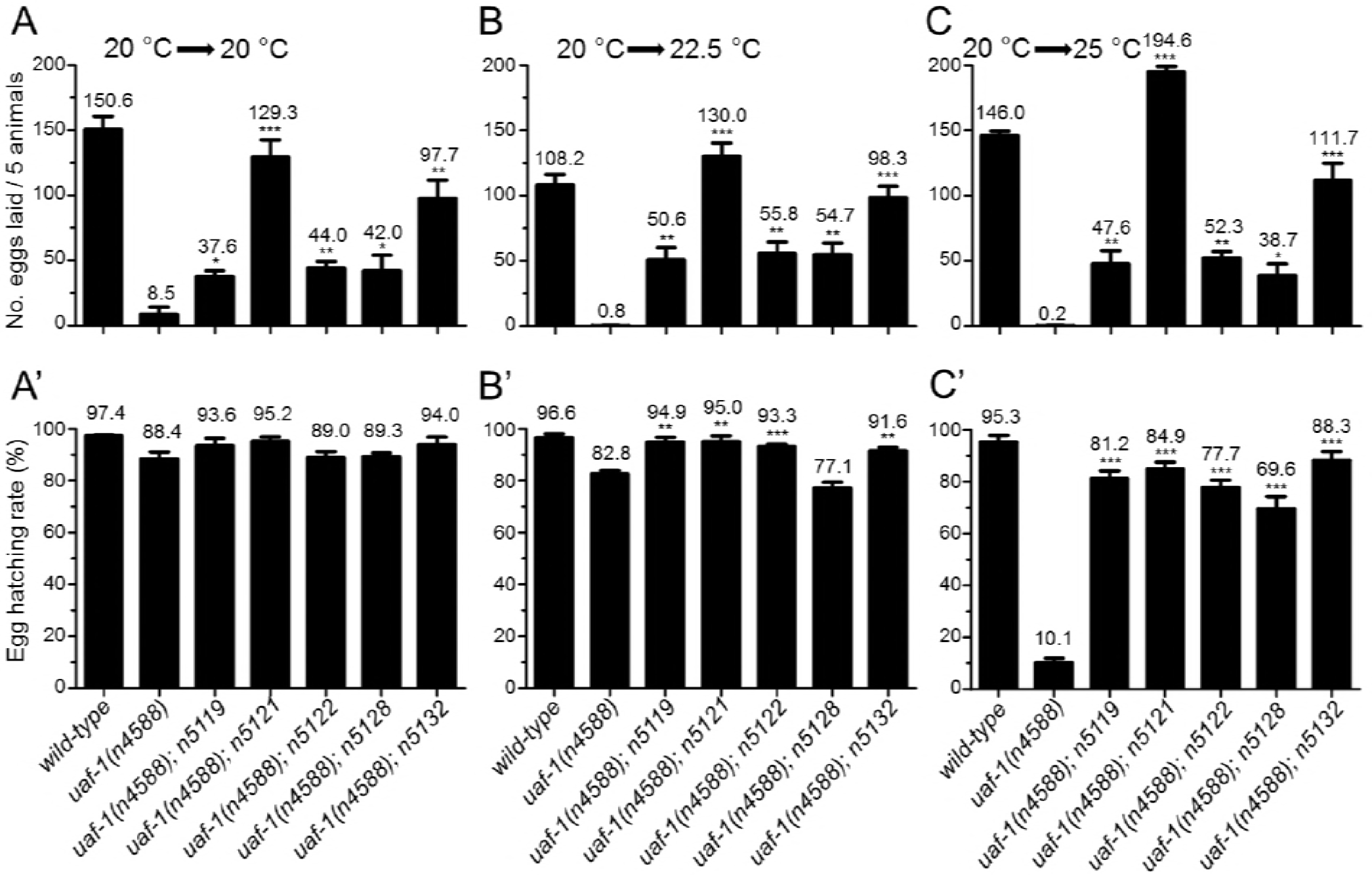
*n5119*, *n5121*, *n5122*, *n5128* and *n5132* can suppress the fertility and egg-hatching defects of *uaf-(n4588*) mutants at high temperatures. Synchronized L4 animals grown at 20 °C were kept at 20 °C or moved to 22.5 °C and 25 °C. Eggs laid in 24 hours were counted and the hatching rates of the eggs 24 hrs after were quantified. (A, A’) At 20 °C, *uaf-1(n4588)* mutants laid significantly fewer eggs, which was significantly improved in five double mutants. Most eggs laid by *uaf-1(n4588)* single mutants or the double mutants could hatch. (B, B’) Egg-laying and egg-hatching rates at 22.5 °C. The phenotypes are similar to those at 20 °C. (C, C’) Eggs-laying and egg-hatching rates at 25 °C. The total numbers of eggs laid by five animals (A, B, C) and the hatching rates of these eggs (A’, B’, C’) are indicated on top of each column. Genotypes are indicated at the bottom. Results were based on 3 to 6 biological replicates. Because *uaf-1(n4588)* mutants lay few eggs, we used a massive number of mutants to generate enough eggs for the egg-hatching analysis in A’ B’ and C’. Statistics: paired two-tailed Student’s t-test. *: p<0.05; **: p<0.01; ***: p<0.001.

Using visible genetic markers, we mapped *n5119* and *n5132* to Chr. I, *n5121* and *n5128* to Chr. III, and *n5122* to Chr. V. *n5119* might be a dominant suppressor, since both *n5119/+* and *n5119/n5119* could suppress the ts-lethality of *uaf-1(n4588)* at 22.5 °C. *n5132* appeared to be a recessive suppressor because *n5132/n5132* could suppress the ts-lethality of *uaf-1(n4588)* at 22.5 °C, while *n5132/+* failed to do so. In this study, we analyzed the gene affected by *n5119* and *n5132.*

### Mutations in the <u>R</u>NA-<u>b</u>inding <u>m</u>otif protein 5 (RBM) gene *rbm-5* can suppress the ts-lethality of *uaf-1(n4588)* mutants

To identify genes affected by *n5119* and *n5132*, we used whole-genome sequencing to survey the coding sequences and splice sites of genes in the two mutants (see Materials and Methods). From the filtered sequence variants, we identified 12 candidate genes for *n5119* and 5 for *n5132* on Chr. I. Interestingly the RNA-binding motif protein gene *rbm-5 (T08B2.5)* is the only candidate mutated in both *n5119* and *n5132* mutants (Fig. 3A and supplementary Table S1).

**Figure 3.**
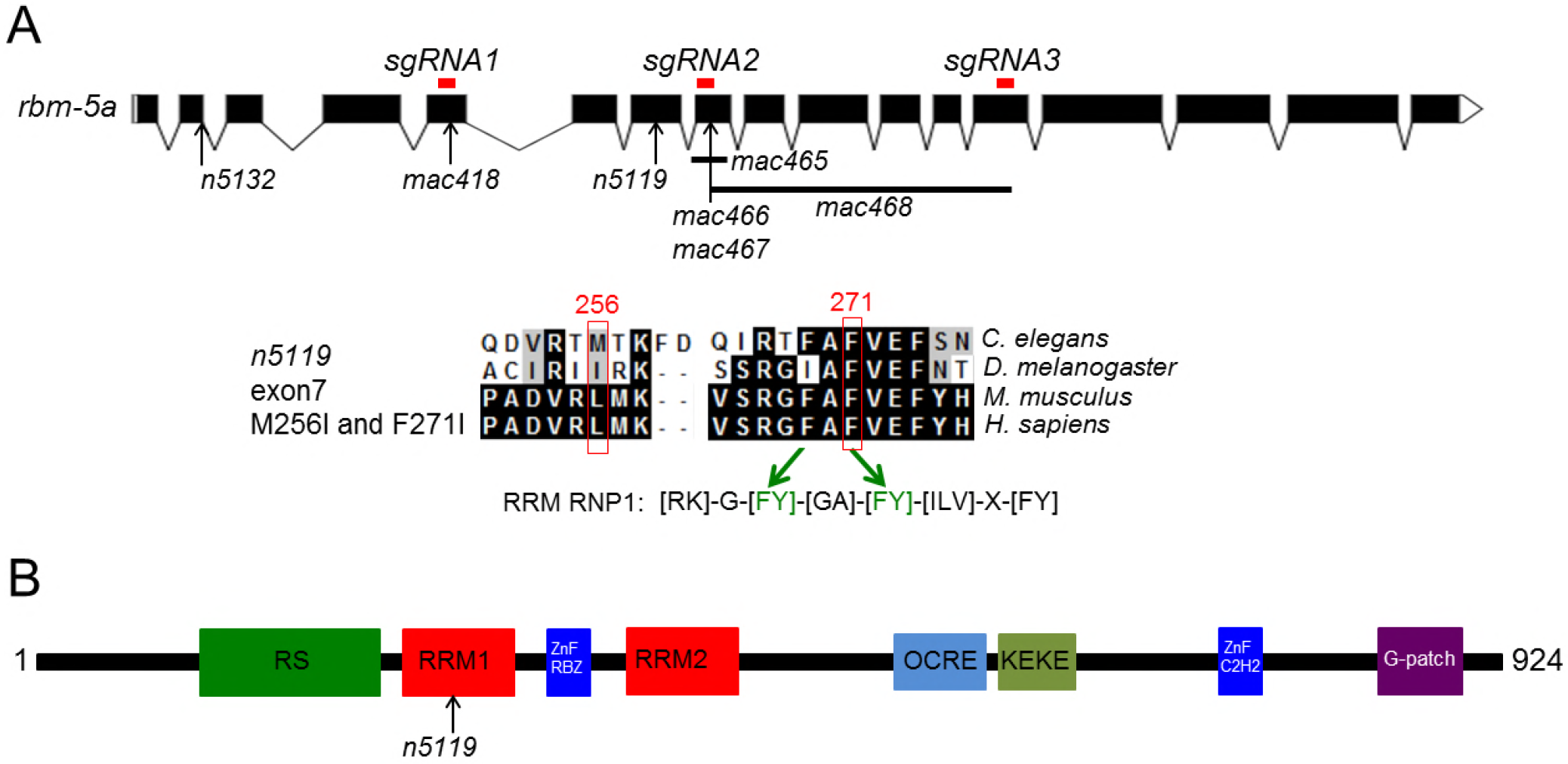
*rbm-5a* gene structure, RBM-5A protein domains and positions of *rbm-5* mutations. (A) Exon-intron structures of the *rbm-5a* isoform (designed using the Exon-Intron Graphic Maker software at www.wormweb.org based on gene sequences at www.wormbase.org). Exonic regions targeted by the CRISPR/Cas9 method are labeled on top as red bars. *n5132* and *n5119* mutations are indicated. The conserved RRM RNP1 sequence is shown, in which the F271I mutation corresponds to the second [FY] residue [57, 58]. *mac418* was obtained using *sgRNA1*, *mac465*, *mac466* and *mac467* were obtained using *sgRNA2. mac468* was obtained using *sgRNA2* and *sgRNA3*. (B) RBM-5 protein domains and positions of the *n5119* missense mutation.

*rbm-5* encodes the *C. elegans* homolog of the mammalian *RBM5/Luca-15/H37* [15] (Fig. S2). *RBM5* was frequently deleted or mutated in lung cancers [19] and can affect alternative splicing, cell proliferation and apoptosis [19, 30, 32, 40]. In *C. elegans*, the function of *rbm-5* is unclear. In the WormBase (www.wormbase.org, WS258), nine *rbm-5* transcript isoforms (*rbm-5a*, *rbm-5b*, *rbm-5c*, *rbm-5d*, *rbm-5e*, *rbm-5f.1*, *rbm-5f.2*, *rbm-5g.1* and *rbm-5g.2*) are annotated (Fig. 3A and S3), representing seven distinct encoding isoforms (*rbm-5a, b, c, d, e, f* and *g*) (Fig. 3A and S3). In *n5119* mutants, the less conserved Met256 (L131 in human) and the conserved Phe271 (F144 in human) encoded by *rbm-5a* are changed to isoleucines (Fig. 3A and S2). Met256 and Phe271 are two residues in the first RNA recognition motif of RBM-5A (Fig. 3B and S2), implying that the changes might affect RNA binding by the RBM-5 protein. In *n5132* mutants, the 5’ splice site in *rbm-5a* intron 2 was mutated from gt to at (Fig. 3A and S3). This mutation is predicted to affect the splicing of at least five isoforms (*rbm-5a, b, d, e* and *f.1)* (Fig. 3A and S3).

To verify that *rbm-5* mutations can suppress the ts-lethality of *uaf-1(n4588)* mutants, we used a modified CRISPR/Cas9 method [41] to generate new mutations in *rbm-5* (Fig. 3A, *sgRNA1*, *sgRNA2*, *sgRNA3*). By this approach, we obtained five deletion mutations in *rbm-5* (Fig. 3A and Table S2). *mac418* and *mac467*, both 4-bp deletions, affect exons 5 and 8 and were predicted to cause frameshifts (Fig. 3A and Table S2). The *mac465* deletion (98 bp) spans from intron 7 and exon8 and was predicted to disrupt the splicing and coding of *rbm-5a* (Fig. 3A and Table S2). *mac466* is a tri-nucleotide deletion predicted to cause a single amino acid deletion in RRM1 (Fig. 3A and Table S2). *mac468* is a 940-bp deletion spanning from exon 8 to exon 13 and was predicted to cause a 238-amino acid deletion if *rbm-5* pre-mRNA was spliced correctly (Fig. 3A and Table S2). Except *mac466*, the other four *mac* alleles *(mac418, mac465, mac467, mac468)* are likely null mutations by disrupting the splicing of *rbm-5* and/or generating truncated RBM-5 proteins (Fig. 3A). We found that homozygous mutants of *mac465* and *mac468* were viable and exhibited no obvious defects (Table 1), suggesting that *rbm-5* is not essential for *C. elegans* survival.

**Table 1:** *rbm-5* mutations can suppress the ts-lethality of *uaf-1(n4588)* mutants at 22.5°C.

| Strains | Survival at 22.5 °C | Survival at 25 °C |
| --- | --- | --- |
| *uaf-1(n4588)* | No | No |
| *rbm-5(n5119)* | Yes | Yes |
| *rbm-5(n5132)* | Yes | Yes |
| *rbm-5(mac465)* | Yes | Yes |
| *rbm-5(mac468)* | Yes | Yes |
| *rbm-5(n5119); uaf-1(n4588)* | Yes | No |
| *rbm-5(n5119)/+; uaf-1(n4588)* | Yes | No |
| *rbm-5(n5132); uaf-1(n4588)* | Yes | No |
| *rbm-5(n5132)/+; uaf-1(n4588)* | No | No |
| *rbm-5(mac418); uaf-1(n4588)* | Yes | No |
| *rbm-5(mac465); uaf-1(n4588)* | Yes | No |
| *rbm-5(mac466); uaf-1(n4588)* | No | No |
| *rbm-5(mac467); uaf-1(n4588)* | Yes | No |
| *rbm-5(mac468); uaf-1(n4588)* | Yes | No |
| *uaf-1(n4588 n5127)* | Yes | Yes (sick) |
| *rbm-5(n5119); uaf-1(n4588 n5127)* | Yes | Yes |
| *rbm-5(n5132); uaf-1(n4588 n5127)* | Yes | Yes |

*mac418*, *mac465*, *mac467* and *mac468* all could recessively suppress the ts-lethality of *uaf-1(n4588)* mutants at 22.5 °C (Table 1). *mac466* failed to suppress *uaf-1(n4588)* (Table 1), suggesting that this mutation does not disrupt RBM-5 function severely enough. None of the *mac* alleles could suppress the lethality of *uaf-1(n4588)* mutants at 25 °C (Table 1). We found that *mac418* failed to complement *n5132* in suppressing the ts-lethality of *uaf-1(n4588)* mutants, suggesting that *n5132* also causes a loss of function in *rbm-5*.

*n5119* causes changes in two RBM-5A residues (the less conserved M256 and the conserved F271) (Fig. 3A) and *n5119* heterozygous mutation can suppress *uaf-1(n4588)* (Table 1). We speculate that *n5119* might be an *rbm-5* allele that functions by a dominant-negative or haploinsufficient mechanism. The similarity of *n5119* and *n5132* in affecting gene expression and alternative splicing (see below) suggests that they both impair *rbm-5*.

We previously isolated *uaf-1(n5127)* as an intragenic suppressor of *uaf-1(n4588)* that enables the animals to survive at 25 °C in an unhealthy state [35]. *n5119* and *n5132* can completely suppress the unhealthiness of *uaf-1(n4588 n5127)* mutants at 25 °C (Table 1), suggesting that these mutations can interact with *uaf-1* in an allele-independent manner.

### *rbm-5* functions in neurons to affect the ts-lethality of *uaf-1(n4588)* mutants

Using an 1183-bp *rbm-5* endogenous promoter (see Materials and Methods) to drive GFP expression, we observed fluorescent signals primarily in multiple head neurons, ventral nerve cord, the CAN neuron, an intestinal cell and a muscle cell in the posterior part of the transgenic animals (Fig. 4A, 4B, 4C and 4D). However, we failed to obtain stable transgenic lines in the wild-type, *n5119* or *n5132* background using an *rbm-5a* cDNA transgene under control of this promoter, probably due to the toxicity of the transgene. To overcome this problem, we used the ubiquitous *eft-3* promoter [42] to drive an *rbm-5a cDNA::GFP* fusion transgene and obtained stable transgenic lines. The RBM-5A::GFP fusion protein was exclusively localized in the nuclei of the numerous visible cells (Fig. 4E, 4F), suggesting that RBM-5 is a nuclear protein.

**Figure 4.**
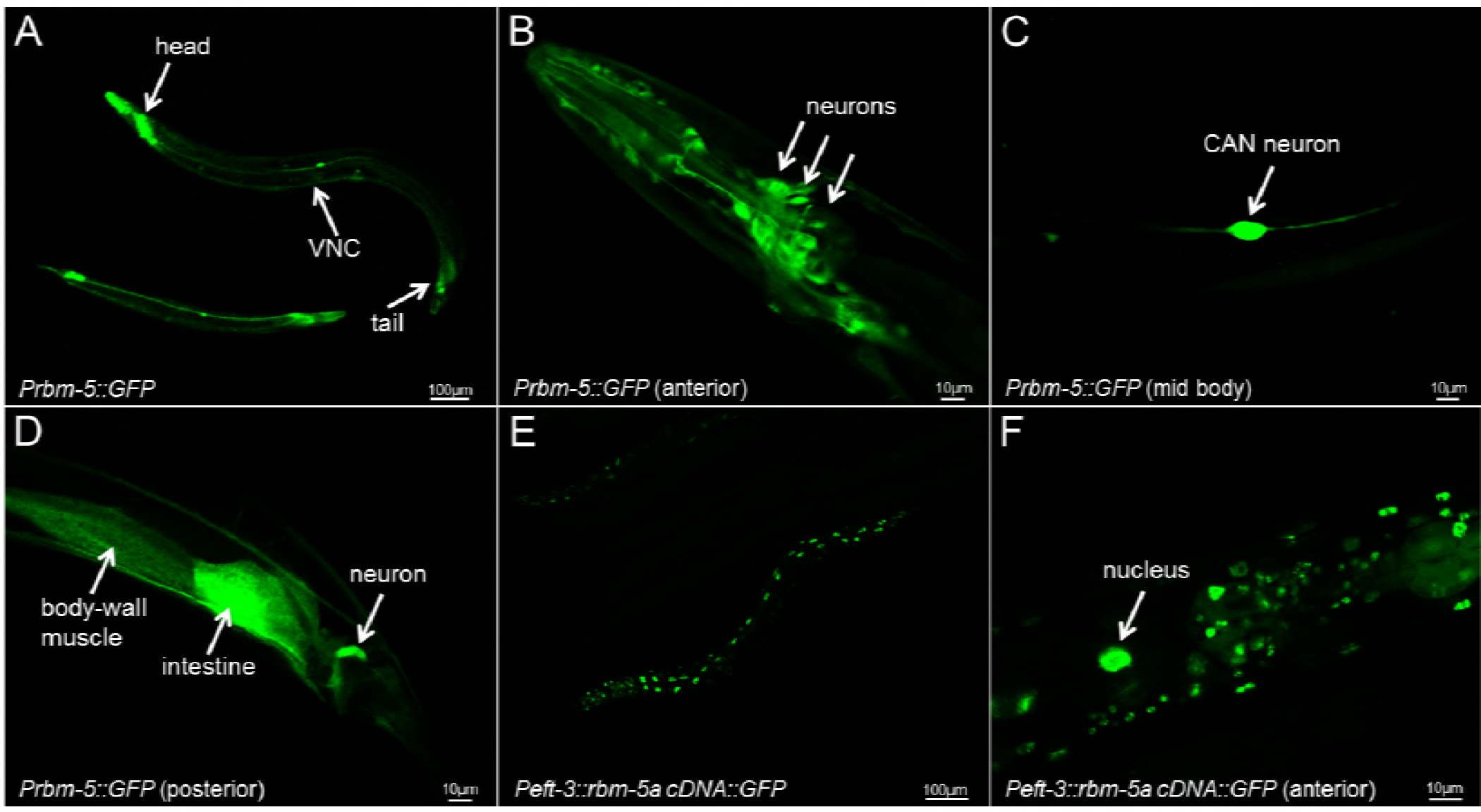
Fluorescent pictures of animals expressing a *Prbm-5::GFP* transcriptional fusion transgene and a *Peft-3::rbm-5a cDNA::GFP* translational fusion transgene. (A) A low resolution picture of *Prbm-5::GFP* transgenic animals showing GFP expression in the anterior, posterior and ventral nerve cord (VNC). High resolution picture of the anterior (B), mid body (C) and posterior (D) of a *Prbm-5::GFP* transgenic animal. Low (E) and high (F) resolution pictures of a *Peft-3::rbm-5a cDNA::GFP* transgenic animal showing that the RBM-5A::GFP fusion protein was exclusively localized in the nuclei.

The *Peft-3::rbm-5a* transgene itself does not affect the survival of animals in different temperatures (Table 2). When placed in the *rbm-5(mac468); uaf-1(n4588)* background, this transgene caused slow growth and reduced fertility at 15 °C, while it led to lethality at 20 °C and 22.5 °C (Table 2). The transgene further caused lethality in *uaf-1(n4588)* single mutant background even at 15 °C (Table 2). These results suggest that the *Peft-3::rbm-5* transgene can rescue the suppression of the *uaf-1(n4588)* ts-lethality by the *rbm-5(mac468)* lf mutation and *rbm-5* overexpression can enhance the lethality caused by *uaf-1(n4588)*.

**Table 2.**
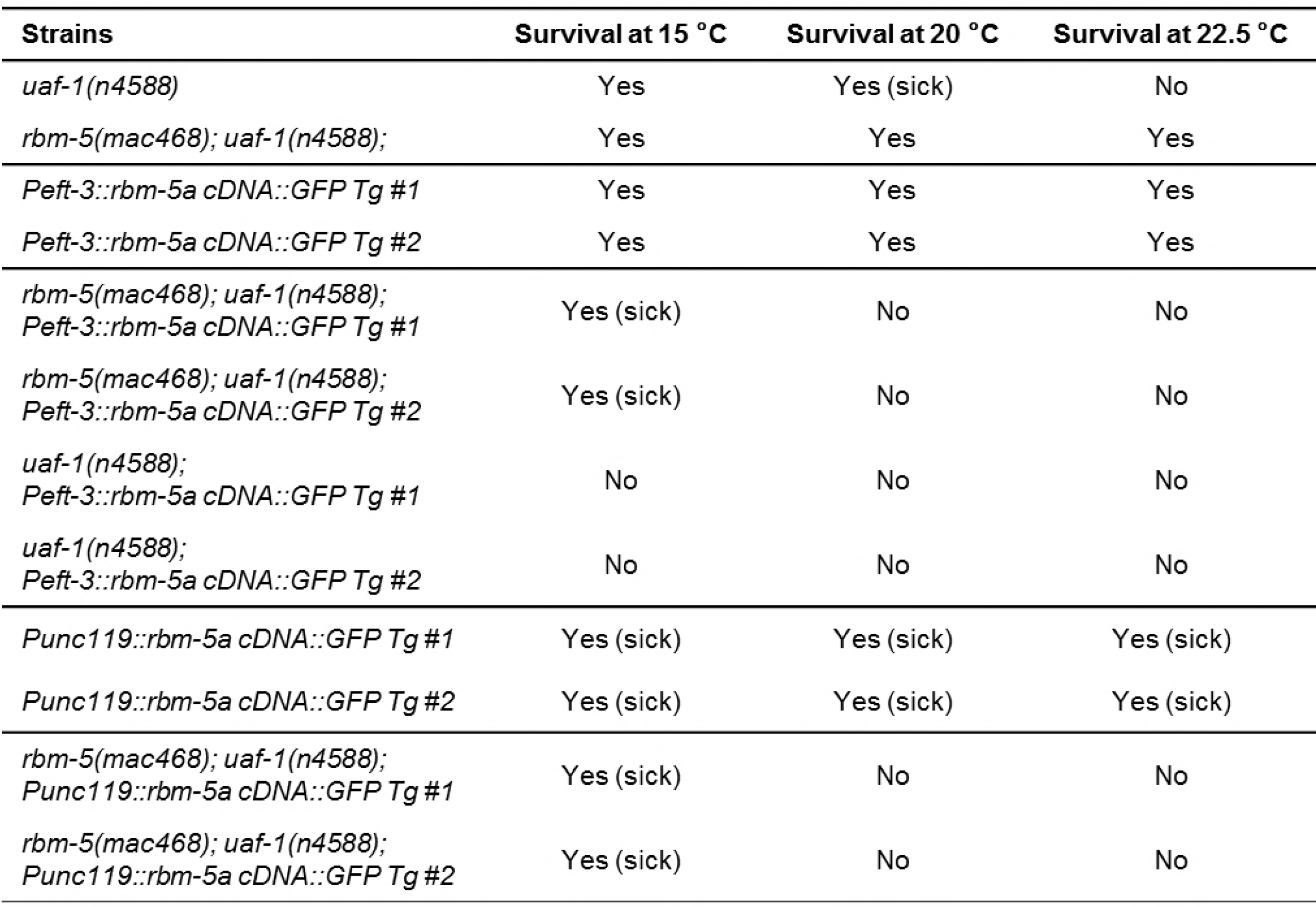
The *rbm-5a cDNA::GFP* transgene under control of the ubiquitous *eft-3* promoter or the neuron-specific *unc-119* promoter could rescue the suppression of the *uaf-1(n4588)* ts-lethality by the *rbm-5(mac468)* mutation.

Since the *rbm-5* endogenous promoter drives reporter expression in multiple neurons (Fig. 4A, 4B, 4C and 4D), we also tested whether neuron-specific expression of *rbm-5* could affect the ts-lethality of *uaf-1(n4588)* mutants by introducing an *rbm-5a* transgene under control of the neuron-specific *unc-119* promoter [43, 44]. In wild-type background, the *Punc-119::rbm-5a cDNA::GFP* transgene caused slow growth and apparently reduced fertility at all temperatures tested (Table 2). When placed in the *rbm-5(mac468); uaf-1(n4588)* background, the transgene led to lethality at at 20 °C and 22.5 °C, similar to the effect of the *Peft-3::rbm-5a* transgene (Table 2). These results suggest that *rbm-5* functions in neurons to promote the deleterious phenotypes caused by *uaf-1(n4588).*

### *rbm-5* mutations could reduce the number of differentially expressed genes and reverse some altered pathways in *uaf-1(n4588)* mutants to wild-type levels

Mammalian RBM5 can affect the splicing of different genes [30–32, 40]. We examined whether *rbm-5* mutations can affect RNA splicing in *C. elegans* and how *rbm-5* interacts with *uaf-1* in doing so.

We performed whole transcriptome shotgun sequencing (RNA-Seq) experiments and compared the embryonic transcriptomes of wild-type, *uaf-1(n4588)* single mutants and *rbm-5(mut); uaf-1(n4588)* double mutants grown at 20 °C. At 20 °C, *uaf-1(n4588)* could produce enough eggs for extracting total RNAs.

We found that *rbm-5* mutations only affected the expression of a small number of genes (3 up, 0 down, 3 total for *n5119*; 9 up, 13 down, 22 total for *n5132*) (Fig. 5A). The number of genes affected was dramatically increased to 194 in *uaf-1(n4588)* embryos (120 up and 74 down) (Fig. 5A). Interestingly, *rbm-5* mutations significantly reduced this number, *e.g., rbm-5(n5119); uaf-1(n4588)* double mutants had 74 genes affected (56 up and 18 down) and *rbm-5(n5132); uaf-1(n4588)* had 52 affected (36 up and 16 down) (Fig. 5A).

**Figure 5.**
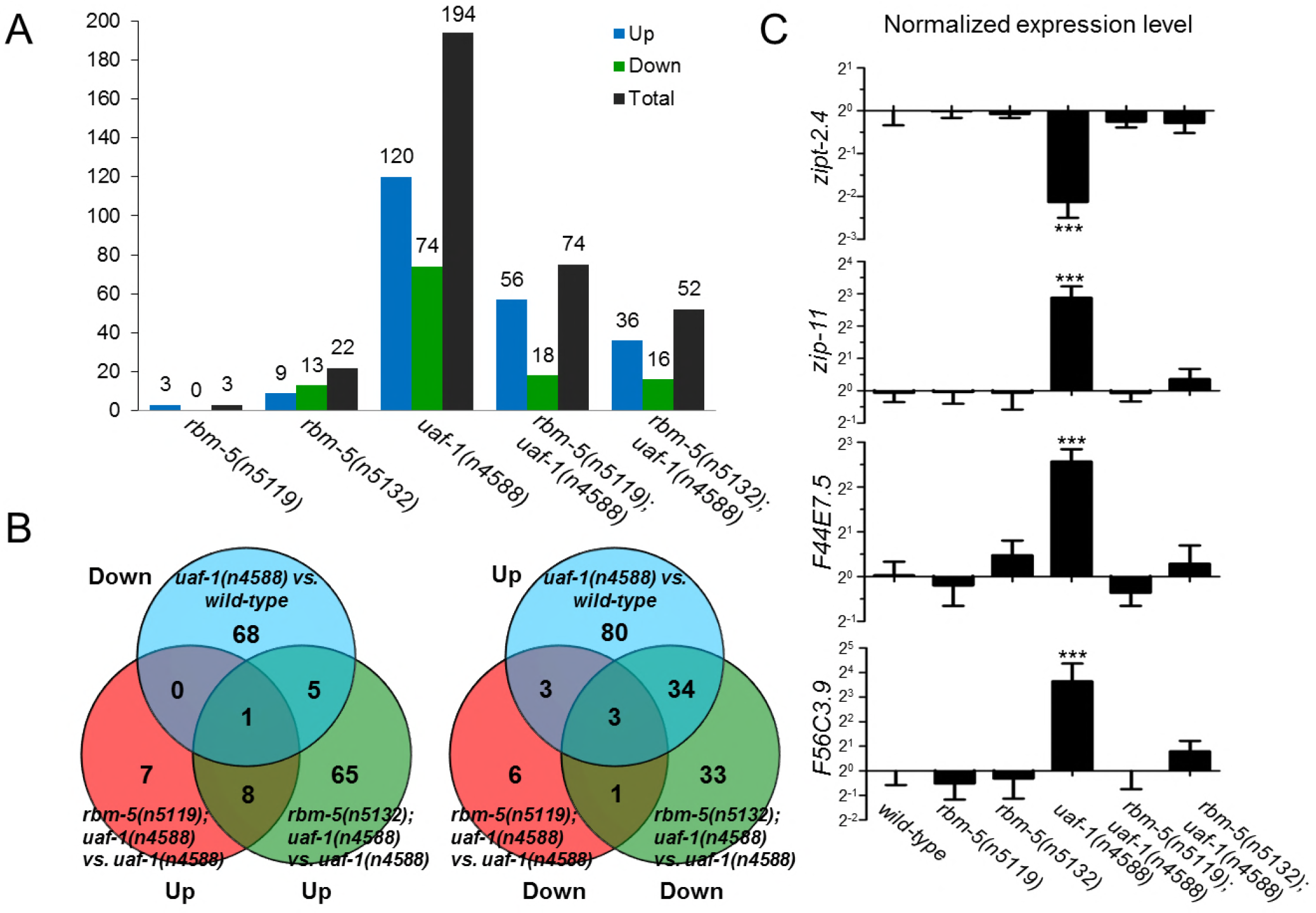
RNA-Seq identified genes differentially expressed in *uaf-1(n4588)* and *rbm-5; uaf-1(n4588)* mutants. (A) Numbers of up-and down-regulated genes in mutants compared to wild type. Genotypes are at bottom. (B) Venn diagram comparison of genes down-regulated in *uaf-1(n4588)* mutants and genes up-regulated in *rbm-5; uaf-1(n4588)* double mutants compared to *uaf-1(n4588)* single mutants, and vice versa. (C) qRT-PCR verification of the expression of four genes predicted by RNA-Seq to be altered by *uaf-1(n4588)* that were reversed by *rbm-5* mutations. Statistics: Bonferroni test with one-way ANOVA. ***: p<0.001.

KEGG (www.genome.jp/kegg/) enrichment analyses of the differentially expressed genes suggest that 40 pathways are affected by *uaf-1(n4588)* (Table S3). These pathways include the metabolism of multiple biomolecules, diseases, synapse functions, signaling pathways and spliceosome, etc, suggesting that *uaf-1* might affect embryonic development by regulating the expression and/or splicing of genes primarily involved in these processes. Compared to wild type, *rbm-5(n5119)* single mutation does not alter any of these pathways (Table S3). *rbm-5(n5132)* single mutation only affects four pathways, including chemical carcinogenesis, cytochrome P450-related metabolism of xenobiotics and drugs, and Parkinson’s disease (Table S3).

Analyses of double mutants suggest that *rbm-5(n5119)* could reverse 35 of the 40 pathways affected by *uaf-1(n4588)* to levels similar to that of wild type. *rbm-5(n5132)* could reverse 16 of the 35 pathways (Table S3). The 16 shared pathways reversed to wild-type levels by both *rbm-5* mutations include nitrogen metabolism, malaria, systemic lupus erythematosus, ubiquinone-related biosynthesis, fat digestion and absorption, glyoxylate and dicarboxylate metabolism, alcoholism, glutamatergic synapse, amino sugar and nucleotide sugar metabolism, arginine and proline metabolism, transcriptional misregulation in cancer, Alzheimer’s disease, PPAR signaling pathway, protein processing in endoplasmic reticulum, peroxisome and spliceosome (Table S3). Though it is unclear how these pathways are involved in *C. elegans* embryonic development, it is plausible that normal expression/splicing of genes in these pathways can sufficiently mitigate the deleterious effects of *uaf-1(n4588)*.

We further looked into the genes differentially expressed (> 2-fold) in *uaf-1(n4588)* and compared that to *rbm-5; uaf-1(n4588)* double mutants (Result S1). Among 194 genes affected by *uaf-1(n4588)*, *rbm-5(n5119)* reversed the expression of 144 (including 78 up-regulated and 66 down-regulated genes) to levels closer to that of wild type, while *rbm-5(n5132)* reversed that of 177 genes (including 111 up-regulated and 66 down-regulated ones) to levels closer to that of wild type (Result S1).

We identified four differentially expressed genes in *uaf-1(n4588)* mutants that were reversed by both *rbm-5* mutations (Fig. 5B), including *zipt-2.4* (down in *n4588), zip-11* (up), *F44E7.5* (up) and *F56C3.9* (up). qRT-PCR verified altered expression of these genes in *uaf-1(n4588)* mutants and the reversion by *rbm-5* mutations as predicted by RNA-seq (Fig. 5C).

### *rbm-5* can modulate the altered splicing caused by *uaf-1(n4588)*

To understand how *rbm-5* interacts with *uaf-1(n4588)* to affect alternative splicing, we used RT-PCR to examine 80 genes alternatively spliced in *uaf-1(n4588)* mutants with the highest statistical significance predicted by RNA-Seq (Table S4). Among these genes, only three (*flp-1*, *mbk-2*, *zipt-2.4*) were verified (Fig. 6A). The splicing of these genes represents three distinct patterns: alternative 3’ SS (*flp-1*), exon skipping (*mbk-2*), and alternative 5’ SS (*zipt-2.4*) (Fig. 6A). *rbm-5* mutations could apparently suppress the altered splicing of *mbk-2* and *zipt-2.4* in *uaf-1(n4588)* mutants (Fig. 6A), while the altered splicing of *flp-1* was not suppressed (Fig. 6A). We had shown that the expression of *zipt-2.4* was significantly decreased in *uaf-1(n4588)* mutants, which was suppressed by *rbm-5* mutations (Fig. 5C). However, the expression levels of *flp-1* and *mbk-2* were not affected by *uaf-1(n4588)* single or *rbm-5; uaf-1(n4588)* double mutations (Fig. S4).

**Figure 6.**
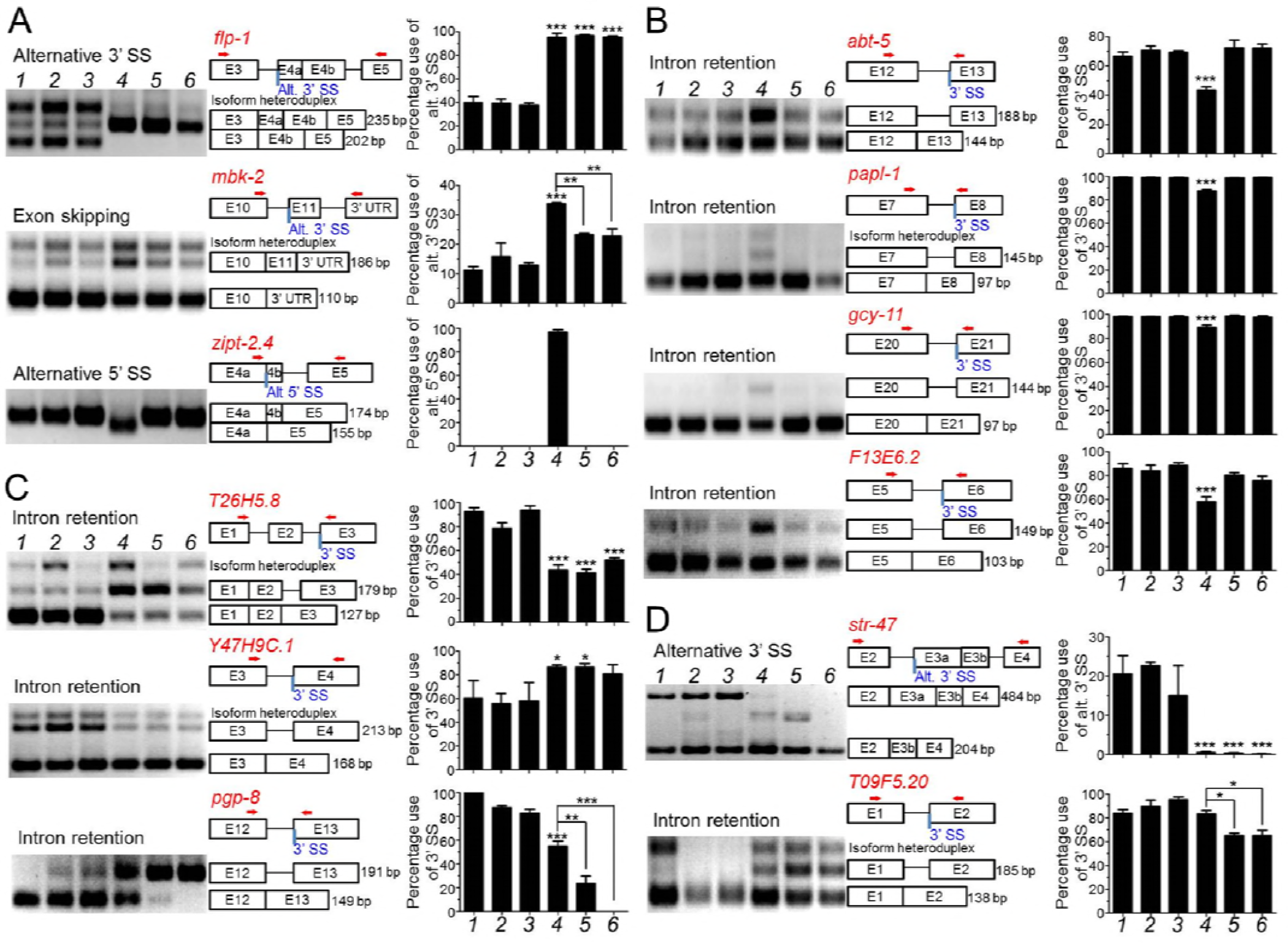
RT-PCR analyses of genes with altered splicing in *uaf-1(n4588)* single and/or *rbm-5; uaf-1(n4588)* double mutants. (A) Three genes with altered splicing in *uaf-1(n4588)* as predicted by RNA-Seq are verified. (B) The altered splicing of four genes with the “gtag” 3’SS in *uaf-1(n4588)* mutants can be suppressed by *rbm-5* mutations. (C) The altered splicing of two genes (*T26H5.8* and *Y47H9C.1*) with the “gtag” 3’SS in *uaf-1(n4588)* mutants was not affected by *rbm-5* mutations and that of *pgp-8* was enhanced by *rbm-5* mutations. (D) For genes with the “ttag” 3’ SS, *rbm-5* mutations did not affect the altered splicing of a gene (*str-47*) caused by *uaf-1(n4588)* while functioned synthetically with *uaf-1(n4588)* to cause altered splicing of another gene (*T09F5.20*). From left to right, genotypes are shown in numerical order: wild type (1), *rbm-5(n5119)* (2), *rbm-5 (n5132* (3), *uaf-1(n4588)* (4), *rbm-5(n5119); uaf-1(n4588)* (5) and *rbm-5(n5132); uaf-1(n4588)* (6). Exon-intron structures and lengths of PCR fragments are indicated. Red arrows: PCR primers. Quantifications are based on three biological replicates. Statistics: Bonferroni test with one-way ANOVA. *: p<0.05; **: p<0.01; ***: p<0.001.

We previously found that *uaf-1(n4588)* could increase the recognition of the “gc<u>ag</u>” 3’ SS and the “gt<u>ag</u>” 3’ SS (conserved splice acceptor underlined) in the *C. elegans unc-93(e1500)* pre-mRNA and *tos-1* pre-mRNA [35, 38], respectively, in which the “g” nucleotide at position −4 might play a critical role for the increased recognition [35]. We also found that the splicing of strong 3’ SS in these genes was not affected by *uaf-1(n4588)* [35, 38]. In *mbk-2*, the 3’ SS of intron 10 is gtag. *uaf-1(n4588)* could increase the recognition of this 3’ SS (Fig. 6A).

To understand how “gtag” affects alternative splicing in other genes, we examined 20 genes containing this sequence as 3’ SS that were predicted by RNA-Seq to be alternatively spliced in *uaf-1(n4588)* mutants (Table S4). The splicing of seven genes, including *abt-5*, *papl-1*, *gcy-11*, *F13E6.2*, *T26H5.8*, *Y47H9C.1* and *pgp-8*, was verified to be altered by *uaf-1(n4588)* (Fig. 6B and 6C). In six genes (*abt-5*, *papl-1*, *gcy-11*, *F13E6.2*, *T26H5.8* and *pgp-8*), the altered splicing was caused by a decreased recognition of the “gt<u>ag</u>” 3’ SS (Fig. 6B and 6C, Table 3). In one gene (*Y47H9C.1*), the altered splicing was caused by an increased recognition of this 3’ SS (Fig. 6C and Table 3). *rbm-5* mutations exerted different effects on these splicing events: they suppressed the decreased recognition of this 3’ SS in four genes (*abt-5*, *papl-1*, *gcy-11*, *F13E6.2*) (Fig. 6B, Table 3), did not obviously affect the altered recognition in two genes (*T26H5.8*, *Y47H9C.1*) and enhanced the decreased recognition in one gene (*pgp-8*) (Fig. 6C, Table 3). Therefore, *uaf-1(n4588)* can affect the recognition of the “gtag” 3’ SS in variable manners and *rbm-5* mutations could suppress or enhance the effects of *uaf-1(n4588)* in a gene-specific manner.

**Table 3.**
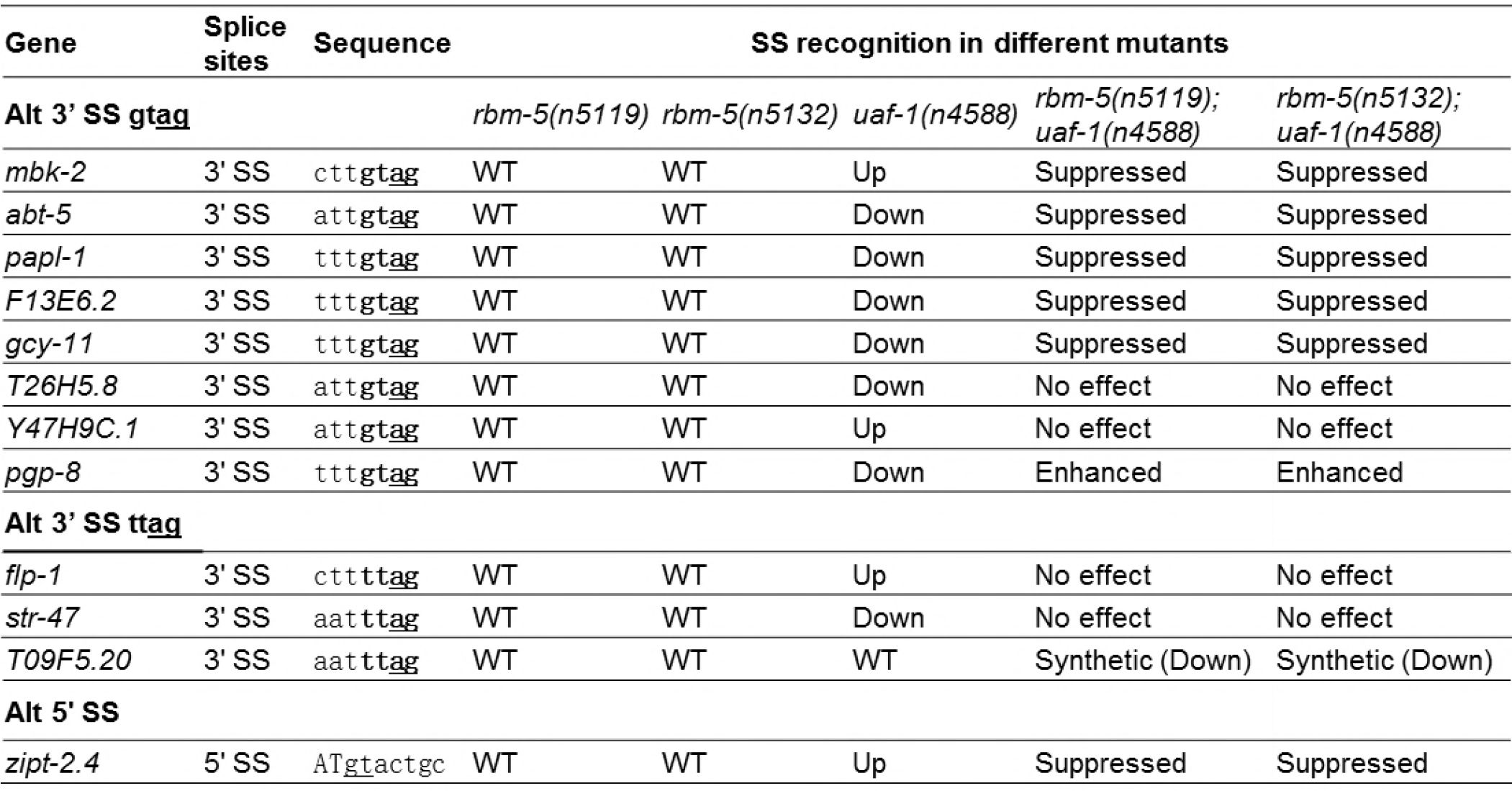
Effects of *rbm-5* mutations on the recognition of SS altered by *uaf-1(n4588).*

| Gene | Splice sites | Sequence | SS recognition in different mutants |  |  |  |  |
| --- | --- | --- | --- | --- | --- | --- | --- |
| **Alt 3' SS gtag** |  |  | *rbm-5(n5119)* | *rbm-5(n5132)* | *uaf-1(n4588)* | *rbm-5(n5119); uaf-1(n4588)* | *rbm-5(n5132); uaf-1(n4588)* |
| *mbk-2* | 3' SS | cttgtag | WT | WT | Up | Suppressed | Suppressed |
| *abt-5* | 3' SS | attgtag | WT | WT | Down | Suppressed | Suppressed |
| *papl-1* | 3' SS | tttgtag | WT | WT | Down | Suppressed | Suppressed |
| *F13E6.2* | 3' SS | tttgtag | WT | WT | Down | Suppressed | Suppressed |
| *gcy-11* | 3' SS | tttgtag | WT | WT | Down | Suppressed | Suppressed |
| *T26H5.8* | 3' SS | attgtag | WT | WT | Down | No effect | No effect |
| *Y47H9C.1* | 3' SS | attgtag | WT | WT | Up | No effect | No effect |
| *pgp-8* | 3' SS | tttgtag | WT | WT | Down | Enhanced | Enhanced |
| **Alt 3' SS ttag** |  |  |  |  |  |  |  |
| *flp-1* | 3' SS | cttttag | WT | WT | Up | No effect | No effect |
| *str-47* | 3' SS | aatttag | WT | WT | Down | No effect | No effect |
| *T09F5.20* | 3' SS | aatttag | WT | WT | WT | Synthetic (Down) | Synthetic (Down) |
| **Alt 5' SS** |  |  |  |  |  |  |  |
| *zipt-2.4* | 5' SS | ATgtactgc | WT | WT | Up | Suppressed | Suppressed |

“tt<u>ag</u>” is the 3’ SS in intron 3 of the *flp-1* gene that was alternatively recognized in *uaf-1(n4588)* mutants (Fig. 6A and Table 3). “tt<u>ag</u>” is more frequently found in the *C. elegans* genome than “gt<u>ag</u>” [45, 46]. We used RT-PCR to examine 20 genes with “tt<u>ag</u>” 3’ SS that RNA-Seq predicted to be alternatively spliced in *uaf-1(n4588)* mutants (Table S4). Only the splicing of one gene *(str-47)* was verified to be altered in *uaf-1(n4588)* mutants due to a decreased recognition of this 3’ SS (Fig. 6D and Table 3). *rbm-5* mutations did not further affect this altered recognition (Fig. 6D and Table 3). The splicing of another gene, *T09F5.20*, was not altered by *uaf-1(n4588)* per se (Fig. 6D). However, it was altered in *rbm-5; uaf-1(n4588)* double mutants as a result of a decreased recognition of the 3’ SS, suggesting a synthetic interaction of *rbm-5* and *uaf-1* (Fig. 6D and Table 3).

We found that the expression levels of five genes, including *gcy-11*, *F13E6.2*, *T26H5.8*, *pgp-8* and *T09F5.20*, were not affected by any single or double mutations (Fig. S4). The expression levels of *abt-5* and *papl-1* were significantly decreased in *uaf-1(n4588)* mutants, while that of *Y47H9C.1* was significantly increased in *uaf-1(n4588)* single or *rbm-5; uaf-1(n4588)* double mutants (Fig. S4). These results suggest that there is no apparent correlation between altered splicing and expression levels in these genes.

It is intriguing that *uaf-1(n4588)* led to the recognition of a cryptic 5’ SS in the *zipt-2.4* gene, which was suppressed by *rbm-5* mutations (Fig. 6A). Mammalian RBM5, RBM6 and RBM10 can bind exonic sequences close to 5’ SS of numerous genes [40], raising a question whether the altered recognition at this cryptic 5’ SS might reflect an *in vivo* function of *rbm-5* and *uaf-1* in regulating splicing at 5’ SS. Alternatively, such a change might result from *uaf-1(n4588)*-induced secondary effect, *e.g.*, the altered splicing of a UAF-1 target gene that was required for the recognition of 5’ SS.

## Discussion

In this study, we identified mutations in the RNA-binding motif gene *rbm-5* as suppressors of the ts-lethality caused by *uaf-1(n4588). rbm-5* interacts with *uaf-1* to affect gene expression and alternative splicing in *C. elegans.* We suggest that the tumor suppressor function of human RBM5 might involve splicing of genes regulated by the U2AF large subunit.

### *C. elegans uaf-1* mutations revealed novel features of the U2AF large subunit

We previously isolated splicing factor gene mutations, *uaf-1(n4588)*, *sfa-1(n4562)* and *mfap-1(n4564 n5214)* [35, 36], as suppressors of the “rubberband” Unc phenotype of *unc-93(e1500gf)* animals. These genes affect alternative splicing in a similar manner [35, 36, 38], implying a functional interaction *in vivo*.

*uaf-1(n4588)* is predicted to change the conserved Thr180 to isoleucine in the UAF-1 protein [35]. Four intragenic suppressive mutations of *uaf-1(n4588)* (M157I, P177L, V179M and T180F) [35] exhibited differential effects on the recognition of 3’ SS *in vivo* [35, 38]. In mammalian U2AF large subunit, the corresponding residues of the four amino acids are M110, S142, M144 and T145, respectively [35, 47]. M110 is a conserved residue in the UHM-ligand motif (ULM) [48], which is also known as the “U2AF small subunit-interacting domain” [35, 49]. S142, M144 and T145 are located in a small region between ULM and RRM1 of the U2AF large subunit, the function of which is unclear.

Recently, a structure-function study of U2AF large subunit suggests that residues of this region (a.a. 141 to 147 in mammalian U2AF large subunit) [50] are important for high-affinity binding to the polypyrimidine tract. A Q147A change could reduce the affinity of U2AF large subunit for the polypyrimidine tract by 5 folds and this region might coordinate with the C-terminal region of RRM2 and the inter-RRM linker region to affect the affinity and specificity of RNA binding [50]. This study corroborates our findings that residues in the region between ULM and RRM1 are important for 3’ SS recognition and alternative splicing in *C. elegans* [35, 38]. Specifically, the *n4588* T180I mutation can lead to, in a gene-specific manner, increased or decreased recognition of weak 3’ SS with a “g” nucleotide at −4 position, suggesting that this mutation likely causes UAF-1 to acquire altered affinity for the short pyrimidine tract of *C. elegans* 3’ SS.

### *rbm-5* mutations suppress the ts-lethality of *uaf-1(n4588)* mutants probably by correcting the altered expression and/or splicing of multiple genes

We found that *uaf-1(n4588)* led to altered expression and/or splicing of numerous genes in *C. elegans* embryos (Fig. 5 and 6). *rbm-5* mutations can reverse the abnormal expression of multiple genes altered by the *uaf-1(n4588)* mutation. As a consequence, *rbm-5* mutations can modify the numerous biological pathways affected by *uaf-1(n4588)* to levels closer to that of wild type (Table S3).

We identified 11 genes that exhibited altered splicing in *uaf-1(n4588)* mutants (Fig. 6). *rbm-5* mutations did not affect the splicing of any of the 11 genes per se. However, *rbm-5* mutations can suppress the altered splicing of six genes, enhance that of one and do not apparently modulate that of the other four genes in the *uaf-1(n4588)* background. We also identified a gene (*T09F5.20*) that exhibited altered splicing only in *rbm-5; uaf-1(n4588)* double mutants (Fig. 6D) but was not affected by any single mutations.

Five genes that we verified in this study (Table S5: *zipt-2.4*, *F44E7.5*, *mbk-2*, *papl-1* and *F13E6.2*) could affect fertility, embryonic and/or larval development or germ-line formation (www.wormbase.org). We postulate that changes in expression and/or splicing of these genes and other unidentified genes together might lead to phenotypes of *uaf-1(n4588)* mutants, and *rbm-5* mutations suppressed *uaf-1(n4588)* probably by correcting the abnormal expression and/or splicing of these genes.

### *uaf-1(n4588)* provides a sensitized background for analyzing the *in vivo* function of RBM-5 in RNA splicing

We previously found that *uaf-1(n4588)* causes both a loss of function and an altered function in UAF-1 [35, 38]. Considering that U2AF subunits are essential for survival in *C. elegans* and *Drosophila* [35, 51–53], the viability of *uaf-1(n4588)* mutants at permissive temperatures provides a critical reagent for understanding the genetic interactions between *uaf-1* and other splicing factors in animals. The effects of *rbm-5* mutations on alternative splicing were only obvious in the presence of *uaf-1(n4588)* (Fig. 6), suggesting that the sensitized genetic background provided by *uaf-1(n4588)* can be used for studying the *in vivo* functions of other splicing factors or mutations. These splicing factors might fine-tune alternative splicing regulated by key splicing factors but lack apparent effects per se.

In mammals, RBM5 can affect splicing especially at weak 3’ SS [30, 32, 40], which might depend on RBM5 binding to multiple different *cis* RNA sequences [31, 32, 40, 54]. RBM5 could also affect the transition from pre-spliceosomal to spliceosomal complexes [30, 33], a possible mechanistic explanation of how RBM5 affects splicing [55]. In our study, though *rbm-5* did not apparently affect RNA splicing per se, *rbm-5* mutations can suppress or enhance, in a gene-specific manner, the altered recognition on weak 3’ SS caused by *uaf-1(n4588)* (Table 3). Therefore, it is a conserved function of RBM-5 to regulate the splicing at weak 3’ SS. It remains unclear whether *C. elegans* RBM-5 affects splicing by mechanisms similar to that of the mammalian homologs.

Unlike other *rbm-5* lf mutations, *rbm-5(n5119)* suppressed *uaf-1(n4588)* by either a dominant-negative or a haploid-insufficient mechanism. *n5119* carries two missense mutations in the RRM1 domain that change the less conserved M256 and the conserved F271 to isoleucines (Fig. 3A). F271 is comparable to F199 of the human U2AF large subunit RRM1 [56, 57], a key aromatic amino acid essential for RNA binding in most RRMs (Fig. 3A) [57, 58]. An F199A mutation can weaken the interaction of U2AF large subunit RRM1 with polypyrimidine consensus sequence by 9 folds [57]. Hence, it is possible that the *n5119* F271I mutation caused reduced affinity of RBM-5 for RNA targets. Currently, our results could not distinguish the dominant-negative or haploinsufficient nature of this mutation. Further analysis of *rbm-5(n5119)* might provide new insights into the function of RBM-5.

### RBM-5 might interact with UAF-1 by a conserved mechanism to regulate RNA splicing

In mouse, RBM5 is an essential gene for embryonic development and the survival of newborns [25] and required for spermatid differentiation and male fertility probably by affecting spermatid RNA splicing [59]. *C. elegans rbm-5* was not required for survival or fertility. However, it can promote the ts-lethality and sterility of *uaf-1(n4588)* mutants, suggesting that it can function in these biological processes.

Recurrent mutations in several splicing factors, including U2AF1 (U2AF35, U2AF small subunit), SF3B1, SRSF2, ZRSR2 were found to be associated with different cancers [60]. The genetic features of these mutations and mouse model studies suggest that *U2AF1*, *SF3B1* and *SRSF2* might promote cancer formation by gain of function or altered function mechanisms and therefore are potentially proto-oncogenes [60]. Recurrent mutations in *U2AF1* and the *RBM5* paralog *RBM10* were also associated with lung adenocarcinomas [23]. In addition, 220 somatic mutations in *RBM5* have been identified in various cancers as annotated in the Cancer Genome Atlas (TCGA) (www.cancergenome.nih.gov). These mutations include premature stop codons, deletions, splice site changes or intronic/exonic changes that potentially affect splicing, consistent with the idea that *RBM5* might function as a tumor suppressor.

We found that 116 somatic mutations in the U2AF large subunit gene have been identified in multiple cancers as annotated at TCGA, with more than two thirds of them found in seven types of cancers, which include uterine corpus endometrial carcinoma, stomach adenocarcinoma, colon adenocarcinoma, bladder urothelial carcinoma, skin cutaneous melanoma, liver hepatocellular carcinoma and lung squamous cell carcinoma. Interestingly, an M110I mutation corresponding to the *n5127* M157I mutation in *C. elegans* UAF-1 [35] was identified in a case of uterine corpus endometrial carcinoma. Mutations in the U2AF large subunit gene and the splicing factor one gene *(sfa-1)* have also been proposed to be potential cancer drivers based on a comprehensive analysis of the mutation profiles of over 400 splicing factors in various cancers [61]. It remains to be determined whether these mutations contribute to the cancer pathogenesis.

In mammals, RBM5 can interact with the U2AF large subunit [30, 32], the spliceosomal SmN/B/B’ proteins [30, 33] and the DExD/H-box protein DHX15 [62]. How these interactions contribute to the tumor suppressor function of *RBM5* is unclear. That *rbm-5* can modulate UAF-1-dependent RNA splicing provides *in vivo* evidence that the interaction of RBM5 and the U2AF large subunit is conserved. It further suggests that the tumor suppressor function of RBM5 might be related to the U2AF large subunit and potentially the U2AF small subunit (U2AF1), a proto-oncogene. Further analyses of the underlying mechanism should provide new insights into the misregulation of RNA splicing in cancers.

## Materials and Methods

### Strains

*C. elegans* strains were grown at 20 °C unless otherwise indicated. N2 (Bristol) is the reference wild type strain [63]. Strains used in this study include:

CSM600 *rbm-5(n5119) I* (backcrossed 4 times, derived from MT17914)
CSM599 *rbm-5(n5132) I* (backcrossed 4 times, derived from MT17927)
CSM601 *rbm-5(n5119) I; uaf-1(n4588 n5127) III*
CSM602 *rbm-5(n5132) I; uaf-1(n4588 n5127) III*
CSM603 *rbm-5(n5119) I; uaf-1(n4588) III*
CSM604 *rbm-5(n5132) I; uaf-1(n4588) III*
CSM911 *rbm-5(mac465) I*
CSM912 *rbm-5(mac468) I*
CSM725 *rbm-5(mac418) I; uaf-1(n4588) III*
CSM838 *rbm-5(mac465) I; uaf-1(n4588) III*
CSM839 *rbm-5(mac466) I; uaf-1(n4588) III*
CSM840 *rbm-5(mac467) I; uaf-1(n4588) III*
CSM847 *rbm-5(mac468) I; uaf-1(n4588) III*
MT14846 *uaf-1(n4588) III*
MT17922 *uaf-1(n4588 n5127) III*
MT17914 *uaf-1(n4588) III; n5119*
MT17916 *uaf-1(n4588) III; n5121*
MT17917 *uaf-1(n4588) III; n5122*
MT17923 *uaf-1(n4588) III; n5128*
MT17927 *uaf-1(n4588) III; n5132*
CSM949 *macEx523[Prbm-5::GFP]*
CSM841 *macEx448[Peft-3::rbm-5a cDNA::GFP; Pmyo-2::mCherry] #1*
CSM842 *macEx449[Peft-3::rbm-5a cDNA::GFP; Pmyo-2::mCherry] #2*
CSM976 *rbm-5(mac468) I; uaf-1(n4588) III; macEx448[Peft-3::rbm-5a cDNA::GFP; Pmyo-2::mCherry] #1*
CSM977 *rbm-5(mac468) I; uaf-1(n4588) III; macEx449[Peft-3::rbm-5a cDNA::GFP; Pmyo-2::mCherry] #2*
CSM990 *macEx533[Punc-119::rbm-5a cDNA::GFP; Pmyo-2::mCherry] #1*
CSM991 *macEx534[Punc-119::rbm-5a cDNA::GFP; Pmyo-2::mCherry] #2*
CSM992 *rbm-5(mac468) I; uaf-1(n4588) III; macEx533[Punc-119::rbm-5a cDNA::GFP; Pmyo-2::mCherry] #1*
CSM993 *rbm-5(mac468) I; uaf-1(n4588) III; macEx534[Punc-119::rbm-5a cDNA::GFP; Pmyo-2::mCherry] #2*

### Temperature shift experiments

Eggs laid or synchronized animals of various developmental stages grown at 15°C or 20°C were moved to incubators with higher temperatures, including 20°C, 22.5°C and 25°C. The egg hatching rate, larval development and/or egg-laying were quantified.

For egg-hatching, 90~150 eggs laid at 15 or 20 °C were shifted to higher temperatures. Unhatched eggs were counted after 24 hours. For larval development, 90~150 L1 animals grown at 15 °C were shifted to higher temperatures and the numbers of adults were counted after three days. For egg-laying, animals were allowed to develop to various larval stages at 15 or 20 °C and then moved to higher temperatures to grow into mid L4 larvae. 5 or10 L4 animals were picked to new plates and allowed to develop and lay eggs for 24 hrs at higher temperatures. The numbers of eggs were counted.

### Identification of *n5119* and *n5132* using genetic mapping and whole-genome sequencing

*n5119* and *n5132* were mapped to Chr. I using visible genetic markers. Similarly *n5121* and *n5128* were mapped to Chr. III and *n5122* to Chr. V.

Animals were washed from NGM plates and starved for several hours in M9. Genomic DNAs were extracted by proteinase K digestion followed by RNase A treatment and two rounds of phenol-chloroform extraction. Three genomic DNA libraries (380 bp inserts) were constructed by Berry Genomics Co., Ltd (Beijing) using Illumina’s paired-end protocol and paired-end sequencing (100-bp reads) was performed on the Illumina HiSeq 2000. Over 4G clean bases were mapped to the N2 genome (Wormbase release 220) after removal of duplicated reads. SNP calling was performed using Genome Analysis Toolkit (GATK) with the N2 genome as reference. 2,517 (*n5119*), 2,888 (*n5121*), 2602 (*n5122*), 2,650 (*n5128*), and 3,108 (*n5132*) SNPs were detected for these strains. SNPs shared among strains were excluded as they were likely derived from common ancestors. To enhance the stringency for mutation identification, we set the depth of reference base (WT) to be <6 in these mutants. Exon or splice site SNPs with a variant quality greater than 30 were selected. Based on these criteria, we obtained 83 (*n5119*), 107 (*n5121*), 81 (*n5122*), 126 (*n5128*), and 83 (*n5132*) SNPs overall. Since *n5119* and *n5132* were mapped to Chr. I, we postulated that a single gene might be affected by both mutations. Comparing SNP variants on Chr. I (12 SNPs *n5119* and 5 for *n5132*) (Table S1) identified *T08B2.5 (rbm-5)* as the only candidate gene that was mutated in both isolates. The identification of *n5121, n5122* and *n5128* is in progress.

### Generation of *rbm-5* mutations using the CRISPR/Cas9 method

We followed the method by Farboud et al. [41] with minor modifications. Plasmids for microinjection were purified using OMEGA’s Midi Plasmid Purification kit (Omega Bio-tek). The following DNA mixture was injected: 50 ng/μl pPD162 (*Peft-3::Cas9-SV40_NLS)*, 25 ng/μl *PU6::sgRNA*, and 20 ng/μl pPD95_86 (*Pmyo-3::GFP*) plasmid as co-injection marker.

F_1_ animals with GFP signals in body-wall muscles were picked to individual plates and the progeny were analyzed for mutations at or near the target sequences by sequencing.

We tested eight *sgRNAs* that target different *rbm-5* exons (Table S6) and found that three *sgRNAs* (Fig. 3A) were efficient in inducing deletions at or near the target regions (Table S6, red target sequences). Two *sgRNAs* generated four deletions in exons 5 and 8 *(mac418, mac465, mac466* and *mac467) (sgRNA1* and *sgRNA2*, Fig. 3A and Table S2). We co-injected *sgRNA2* and *sgRNA3* to generate large deletions and isolated a 940-bp deletion (*mac468*) that covered a region from RRM1 to the OCRE domain of the RBM-5A protein (Fig. 3A and Table S2)

### Plasmids

To construct the *Prbm-5::GFP* plasmid, a PCR-amplified *rbm-5* promoter (1183 bp upstream of the *rbm-5* start codon) were subcloned to pPD95_79 using *SbfI/XmaI* restriction sites.

To construct the *Peft-3::rbm-5a cDNA::GFP* plasmid, a PCR-amplified *eft-3* promoter (a 597-bp (−16 to −612 base) fragment upstream of the *eft-3* start codon) was subcloned to pPD95_79 using *SbfI/XmaI* sites. The *rbm-5a* cDNA was amplified and subcloned to the *pPD95_79-Peft-3* backbone using *XmaI/KpnI* sites. To construct the *Punc-119::rbm-5a cDNA::GFP* plasmid, a promoter of the neuron-specific *unc-119* gene [43, 44] (2188 bp upstream of the *unc-119a* start codon) was amplified and subcloned to *Peft-3::rbm-5a cDNA::GFP* by replacing *Peft-3* using *SbfI/XmaI* sites. PCR primers are listed in Table S7.

Transgenes were crossed to different mutant backgrounds to test the effects on survival.

### Transgene experiments

Germline transgene experiments were performed as described (Mello et al. 1991). For *Prbm-5::GFP* transcriptional reporter, a transgene solution containing 20 ng/μl *Prbm-5::GFP* was injected to wild type. For *Peft-3::rbm-5a cDNA::GFP* reporter, a mixture containing 1 ng/μl transgene and 2.5 ng/μl pCFJ90 *(Pmyo-2::mCherry)* plasmid as co-injection marker was injected to wild type. For *Punc-119::rbm-5a cDNA::GFP* reporter, a mixture containing 0.2 ng/μl transgene and 2.5 ng/μl pCFJ90 was injected to wild type. We tried but failed to obtain stable lines with a *Prbm-5::rbm-5a cDNA* transgene under control of the 1183-bp *rbm-5* endogenous promoter at concentrations of 0.1, 1, 10 or 50 ng/μl in wild-type or double mutant backgrounds, suggesting that overexpressing *rbm-5* using its endogenous promoter is highly toxic.

Transgenic animals were observed using a Leica TCS SP5 II laser confocal microscope and GFP-positive cells were identified based on the anatomical and morphological characteristics described in WormAtlas (www.wormatlas.org).

### Whole transcriptome shotgun sequencing (RNA-Seq)

Synchronized adult animals were bleached to obtain enough eggs, which were washed three times with H_2_O. Total RNA was prepared using Trizol (Invitrogen), treated with RNase-Free DNase I (New England Biolabs) and incubated at 75 °C for 10 min to inactivate DNase I. RNA-Seq was performed by in-house scientists at Annoroad Gene Technology Corporation (Beijing)

Raw data was processed with Perl scripts to ensure the quality of data used in further analysis. Bowtie2 was used for building the genome index, and Clean Data was mapped to the WBcel235 alignments using TopHat v2.0.12. The Integrative Genomics Viewer was used to analyze the mapping results by the heatmap, histogram, scatter plot or other styles.

Fragments counts were obtained with HTSeq v0.6.0 and WBcel235 Ensembl annotation. Gene expression analysis was performed using DEGSeq 1.18.0. The q-value was assigned to each gene and adjusted by the Benjamini and Hochberg normalization for controlling the false discovery rate (FDR). Genes with *q* ≤ 0.05 and |log2_ratio|≥1 were identified as differentially expressed genes (DEGs). Alternative splicing analysis was performed with Asprofile 1.0.4 and Cuffcompare 2.2.1 by constructing *de novo* annotation from the Ensembl input and merged alignment files. KEGG pathway analysis of differentially expressed genes was carried out using Hypergeometric test.

### RT-PCR experiments

Total RNAs were prepared from eggs collected from bleached adults using Trizol (Invitrogen), treated with RNase-Free DNase I (New England Biolabs) and incubated at 75 °C for 10 min to inactivate DNase I. First-strand cDNA was synthesized with random hexamer oligonucleotides using Maxima First Strand cDNA Synthesis Kit (Thermo Fisher). For alternative splicing, 3 biological replicates of each strain were analyzed and the proportion of each splice isoform was quantified using the NIH ImageJ software.

qRT-PCR was performed on 3 biological replicates of each strain using the Maxima SYBR Green qPCR Master Mix (Thermo Scientific). Fluorescence signals were detected using LightCycler^®^ 96 Instrument (Roche). Each 30 μl PCR reaction contained 1 to 10 ng RT template, 0.5 mM PCR primers and 15 μl 2 x SYBR Green PCR Master Mix. After a preincubation step (95 °C for 10 min), two-step amplification was performed using 40 cycles of denaturation (95 °C for 15 s) and annealing (60 °C for 30 s).

The genotype examined include wild type, *rbm-5(n5119), rbm-5(n5132), uaf-1(n4588), rbm-5(n5119); uaf-1(n4588)* and *rbm-5(n5132); uaf-1(n4588)*. The genes examined include *zipt-2.4*, *zip-11*, *F44E7.5*, *F56C3.9*, *flp-1*, *mbk-2*, *abt-5*, *papl-1*, *gcy-11*, *F13E6.2*, *T26H5.8*, *Y47H9C.1*, *pgp-8* and *T09F5.20*. We used the *crt-1* gene as loading control as its expression was abundant in embryos and its FPKM (fragments per kilobase of gene per million mapped reads) was similar across the strains based on RNA-Seq results. PCR primers for detecting expression levels or alternative splicing are listed in Table S7.

## Data availability

All RNA-Seq datasets generated and analyzed during the current study are available in the GEO (https://www.ncbi.nlm.nih.gov/geo/) with the accession number GSE115695. All other data are available from the corresponding author upon reasonable request.

## Statistic analysis

*P* values were determined by Paired two-tailed Student’s *t*-test or Bonferroni’s multiple comparison test using GraphPad Prism 5.0 software.

## Acknowledgement

This study was initiated in the laboratory of H. Robert Horvitz (HRH). H.R.H. is the David H. Koch Professor of Biology at the Massachusetts Institute of Technology and is an Investigator of the Howard Hughes Medical Institute. We thank members of the Ma laboratory for suggestions. The study is supported by National Natural Science Foundation of China grants (No. 31371253, No. 31571045) and a MOST grant (2016YFC1201805) to LM. Some strains were provided by the CGC, which is funded by NIH Office of Research Infrastructure Programs (P40 OD010440).

## Author Contributions

Conceived and designed the experiments: CZ, XG, LM. Performed the experiments: CZ, XG, SH, WG, JX, LM. Analyzed the data: CZ, YM, LM. Wrote the paper: CZ, YM, LM.

## Supplementary Figure legends and Tables

**Figure S1.**
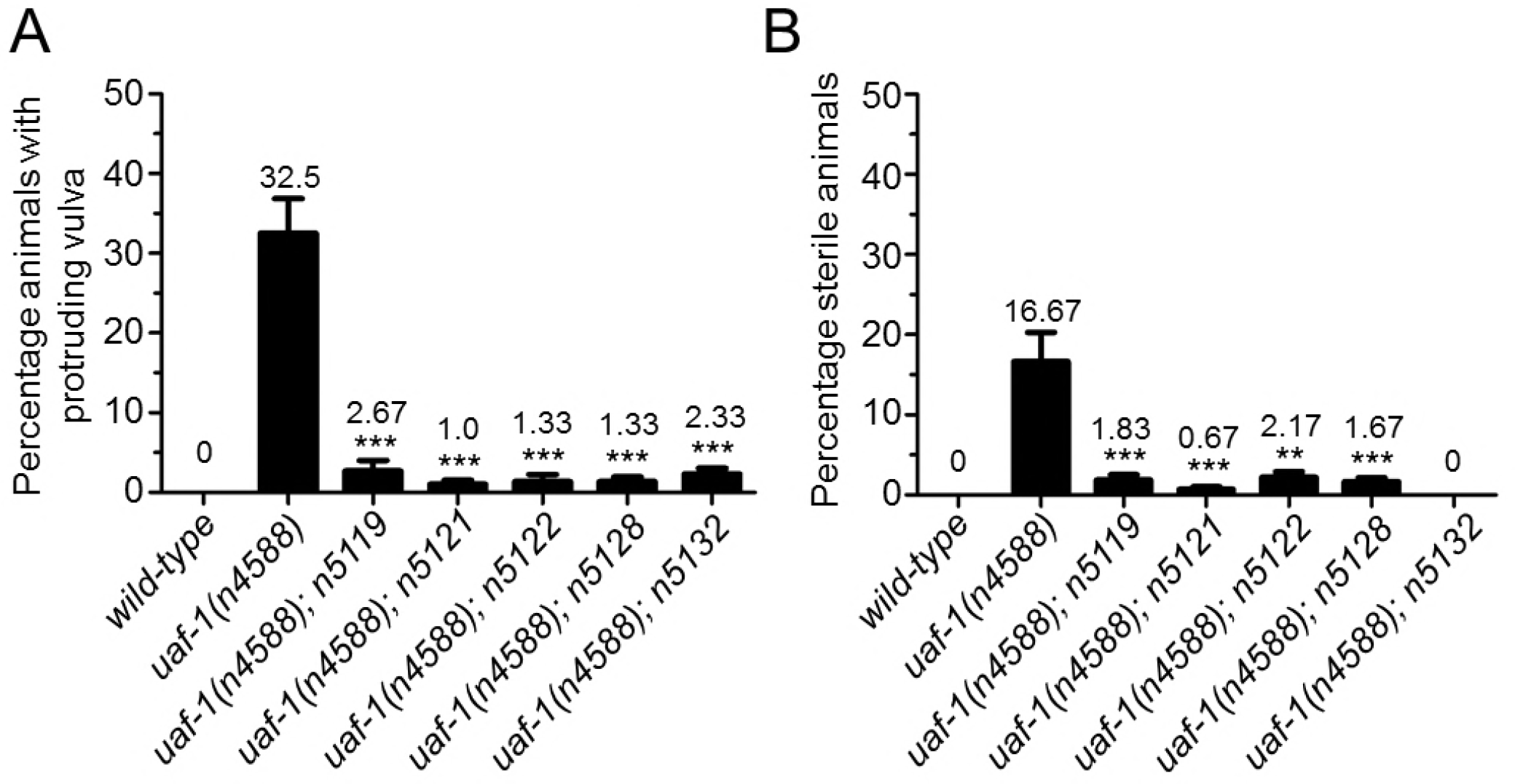
Extragenic mutations can suppress the protruding vulva (Pvl) and sterility (Ste) phenotype of *uaf-1(n4588)* animals. Synchronized L1 animals were grown at 20 °C to young adults and the phenotypes were analyzed. (A) 33% *uaf-1(n4588)* mutants exhibited obvious Pvl phenotype, which was strongly suppressed by the five extragenic mutations. (B) 17% *uaf-1(n4588)* mutants were obviously Ste, which was strongly suppressed by the extragenic mutations. Results were from at least 3 biological replicates and ~100 animals were analyzed in each replicate. Statistics: paired twotailed Student’s t-test.*: p<0.05; **: p<0.01; ***: p<0.001.

**Figure S2.**
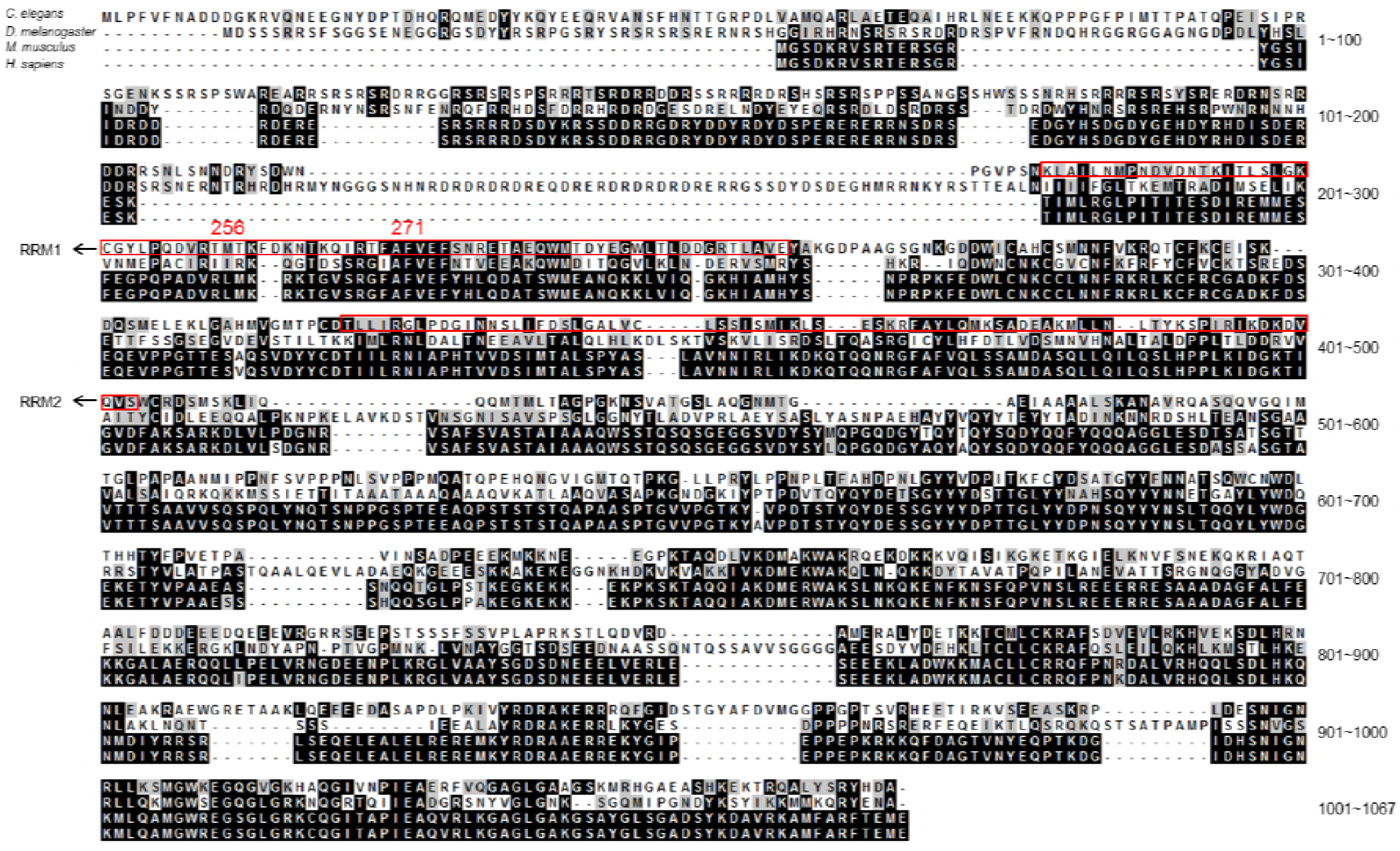
Protein sequence alignment of RBM-5 homologs in *C. elegans*, *Drosophila*, mouse and human. The two amino acid residues mutated in *n5119*, M256 and F271, are indicated. The two RNA recognition motifs, RRM1 from a.a. 224 to 302 and RRM2 from a.a. 363 to 435, are enclosed.

**Figure S3.**
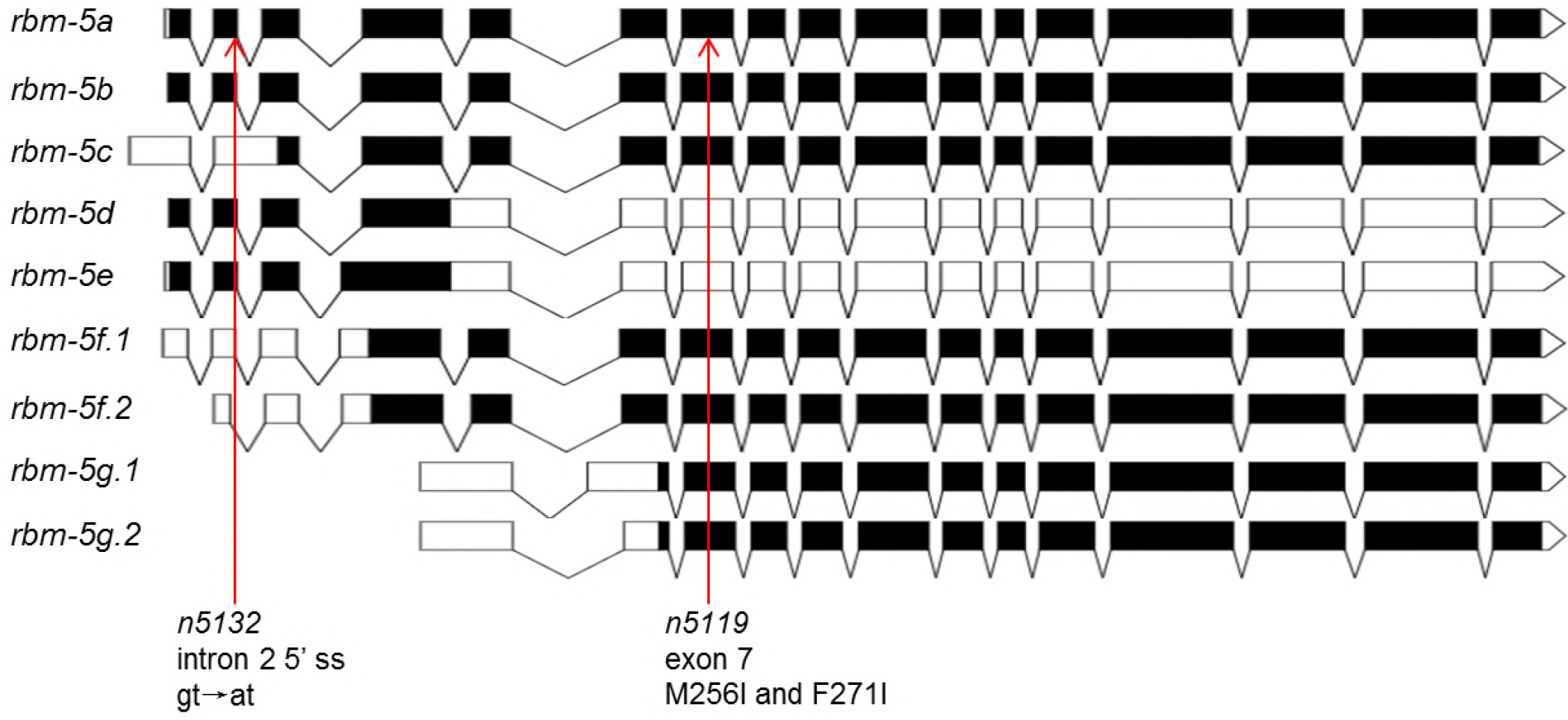
Exon/intron structures of *rbm-5* isoforms. Designed using the Exon-Intron Graphic Maker software at www.wormweb.org based on gene sequence information at www.wormbase.org.

**Figure S4.**
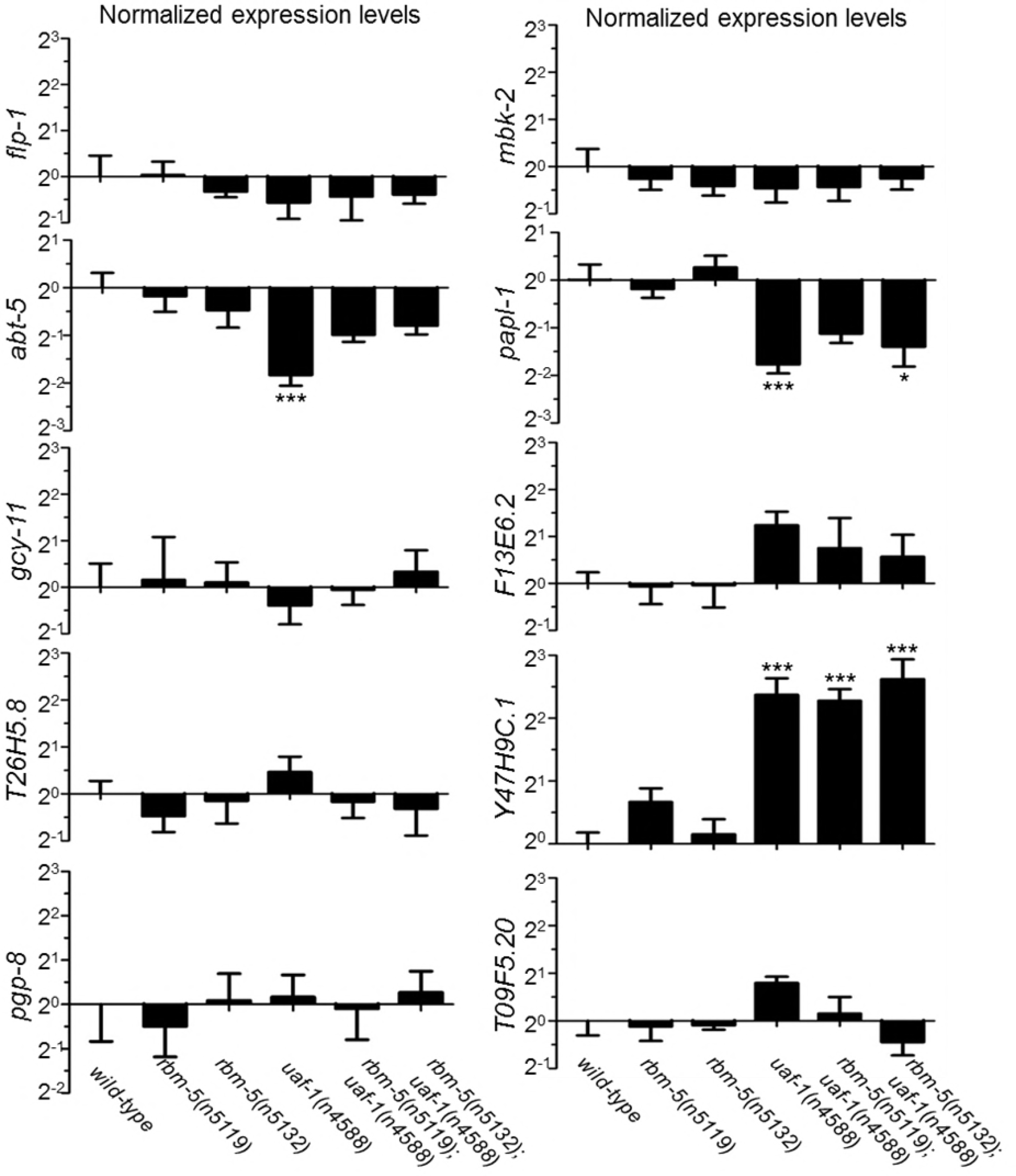
Relative expression levels of ten genes with altered splicing in *uaf-1(n4588)* mutants as shown in Figure 6. Genotypes are labeled at bottom. *str-47* expression level was below the sensitivity of the qRT-PCR experiments and could not be quantified. Statistics: Bonferroni test with one-way ANOVA. *: p<0.05; ***: p<0.001.

**Table S1.** List of candidate genes for *n5119* and *n5132* based on whole-genome sequencing.

| Allele | Candidate genes from whole-genome sequencing |  |  |  |  |  |
| --- | --- | --- | --- | --- | --- | --- |
| *n5119* | *Y23H5B.4* | *Y110A7A.9* | *W05F2.3* | *W02D9.10* | *unc-54* | ***T08B2.5*** |
|  | *T05F1.11* | *mus-101* | *mau-2* | *let-526* | *dyf-1* | *C41G7.3* |
| *n5132* | *nduf-7* | *Y18H1A.11* | *Y106G6H.4* | ***T08B2.5*** | *pad-1* |  |

**Table S2.**
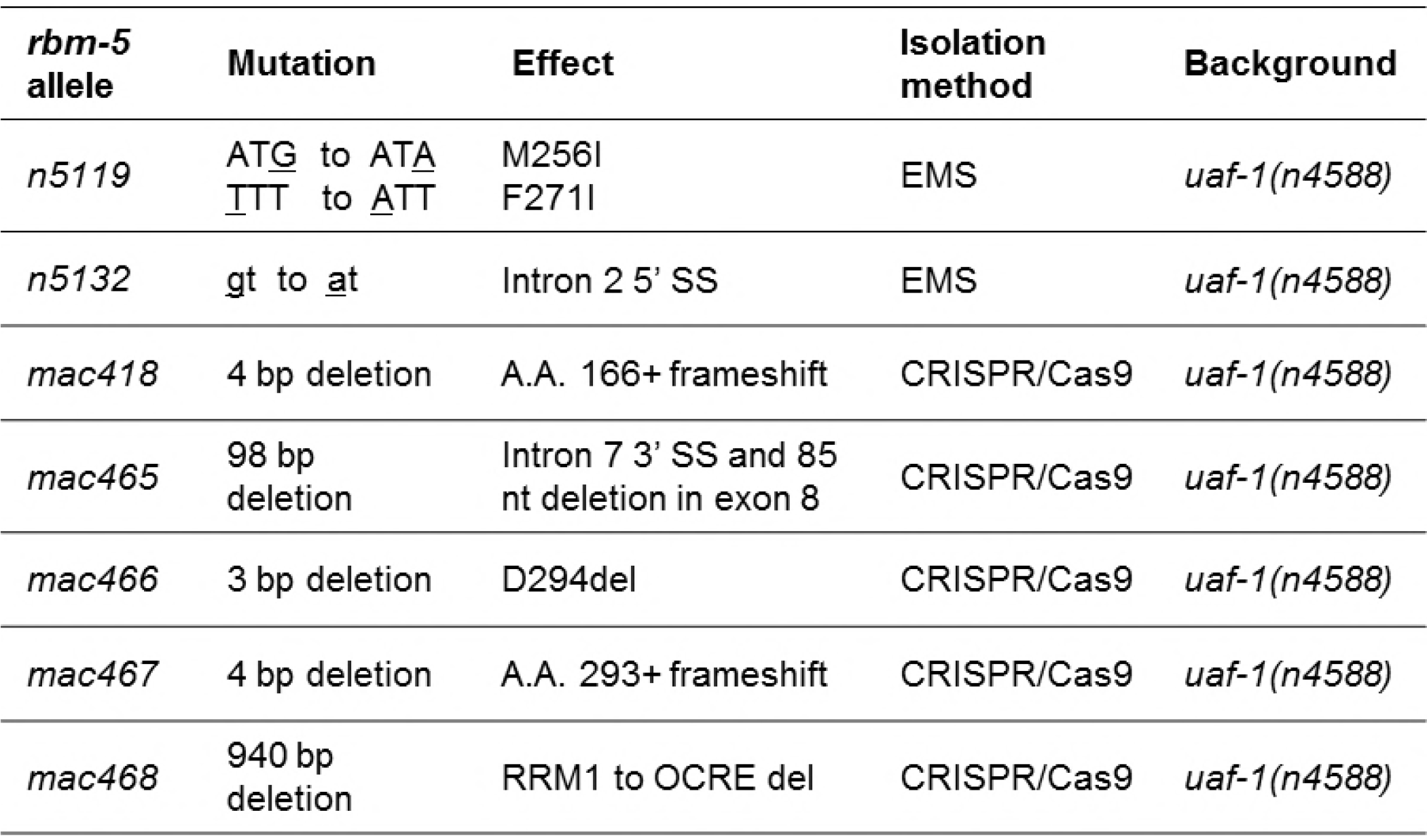
List of *rbm-5* mutations obtained from the EMS mutagenesis screen and the CRISPR/Cas9 method.

**Table S3.** List of 40 pathways altered by *uaf-1(n4588)* based on KEGG analysis and the effects of *rbm-5* mutations on these pathways.

| Pathway | *rbm-5(n5119)* | *rbm-5(n5132)* | *uaf-1(n4588)* | *rbm-5(n5119); uaf-1(n4588)* | *rbm-5(n5132); uaf-1(n4588)* |
| --- | --- | --- | --- | --- | --- |
| Nitrogen metabolism | No effect | No effect | up/down | No effect | No effect |
| Malaria | No effect | No effect | up | No effect | No effect |
| Systemic lupus erythematosus | No effect | No effect | up | No effect | No effect |
| Ubiquinone and other terpenoid-quinone biosynthesis | No effect | No effect | up | No effect | No effect |
| Fat digestion and absorption | No effect | No effect | up | No effect | No effect |
| Glyoxylate and dicarboxylate metabolism | No effect | No effect | up | No effect | No effect |
| Alcoholism | No effect | No effect | up | No effect | No effect |
| Glutamatergic synapse | No effect | No effect | up | No effect | No effect |
| Amino sugar and nucleotide sugar metabolism | No effect | No effect | up | No effect | No effect |
| Arginine and proline metabolism | No effect | No effect | up | No effect | No effect |
| Transcriptional misregulation in cancer | No effect | No effect | up | No effect | No effect |
| Alzheimer's disease | No effect | No effect | up | No effect | No effect |
| PPAR signaling pathway | No effect | No effect | down | No effect | No effect |
| Protein processing in endoplasmic reticulum | No effect | No effect | down | No effect | No effect |
| Peroxisome | No effect | No effect | down | No effect | No effect |
| Spliceosome | No effect | No effect | down | No effect | No effect |
| Alanine, aspartate and glutamate metabolism | No effect | No effect | up/down | No effect | down |
| Glutathione metabolism | No effect | No effect | up/down | No effect | down |
| GABAergic synapse | No effect | No effect | up | No effect | down |
| Calcium signaling pathway | No effect | No effect | up | No effect | up |
| Biosynthesis of amino acids | No effect | No effect | up | No effect | up |
| Cysteine and methionine metabolism | No effect | No effect | up | No effect | up |
| HTLV-I infection | No effect | No effect | up | No effect | up |
| Huntington's disease | No effect | No effect | up | No effect | up |
| Drug metabolism - other enzymes | No effect | No effect | down | No effect | up |
| Ascorbate and aldarate metabolism | No effect | No effect | down | No effect | up |
| Porphyrin and chlorophyll metabolism | No effect | No effect | down | No effect | up |
| Steroid hormone biosynthesis | No effect | No effect | down | No effect | up |
| Pentose and glucuronate interconversions | No effect | No effect | down | No effect | up |
| Starch and sucrose metabolism | No effect | No effect | down | No effect | up |
| Retinol metabolism | No effect | No effect | down | No effect | up |
| Chemical carcinogenesis | No effect | down | up/down | No effect | up/down |
| Metabolism of xenobiotics by cytochrome P450 | No effect | down | up/down | No effect | up/down |
| Drug metabolism - cytochrome P450 | No effect | down | up/down | No effect | up/down |
| Parkinson's disease | No effect | down | up | No effect | up/down |
| Glycine, serine and threonine metabolism | No effect | No effect | up/down | down | up/down |
| Phagosome | No effect | No effect | down | down | up/down |
| Tuberculosis | No effect | No effect | down | down | up/down |
| Histidine metabolism | No effect | No effect | down | down | down |
| Carbon metabolism | No effect | No effect | down | down | down |

**Table S4.**
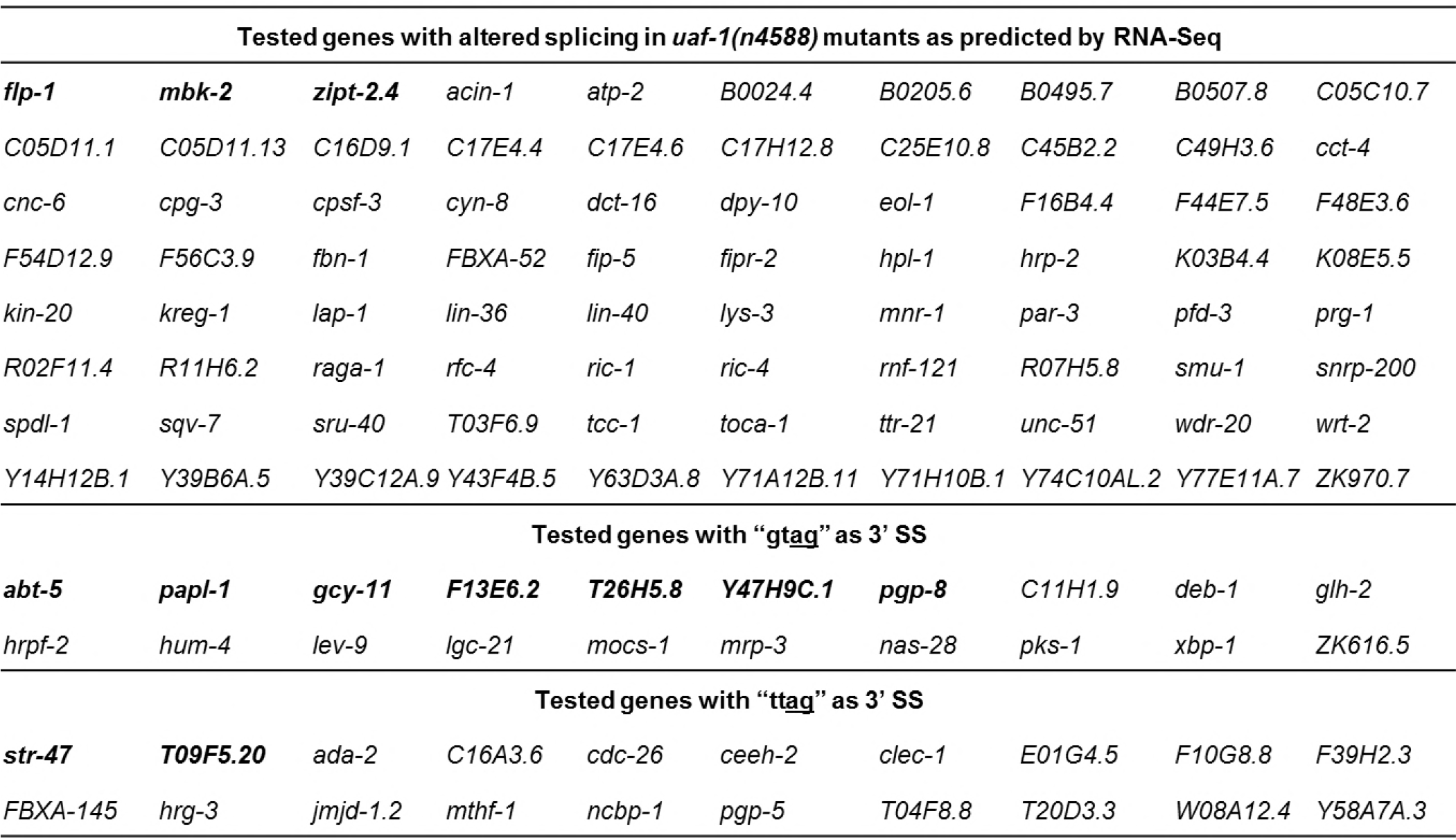
List of genes examined for altered splicing in *uaf-1(n4588)* mutants by RT-PCR. Verified genes are in bold.

**Table S5.**
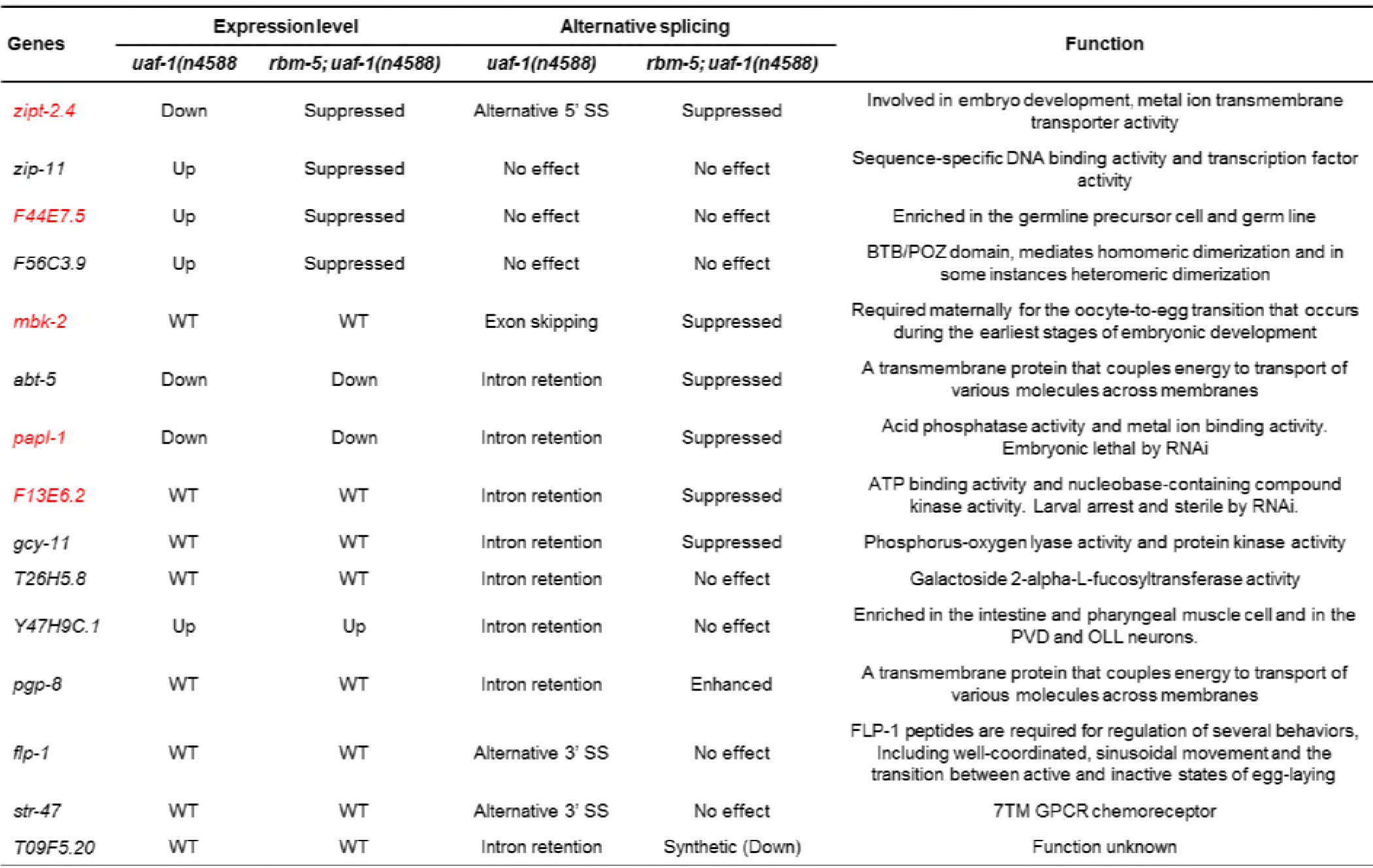
List of genes with verified expression changes and/or altered splicing and their annotated functions (www.wormbase.org).

**Table S6.**
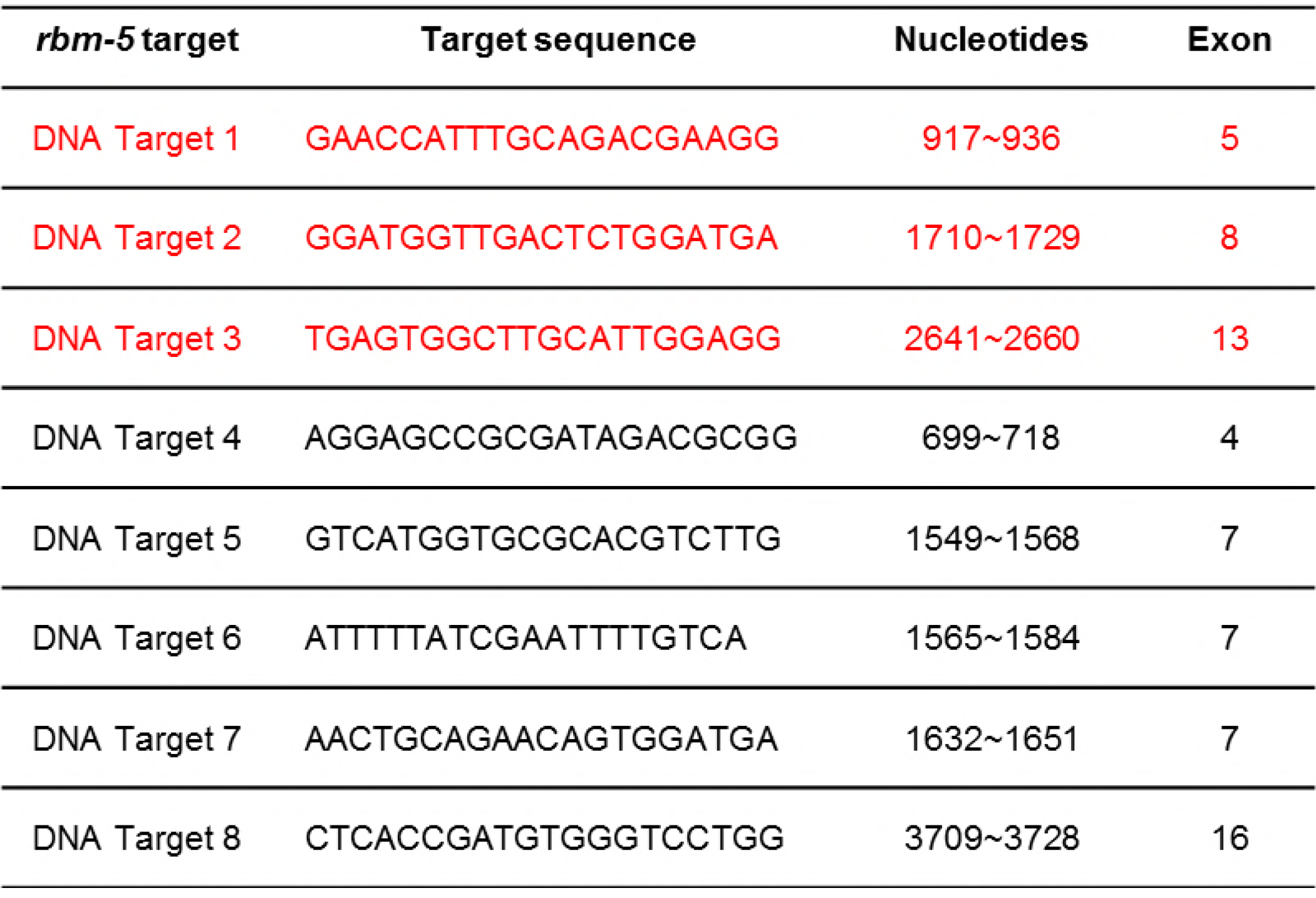
*rbm-5* exonic sequences used as targets in the CRISPR/Cas9 experiment. Effective target sequences are listed on top in red, corresponding to *sgRNA1*, *sgRNA2* and *sgRNA3*, respectively.

**Table S7.**
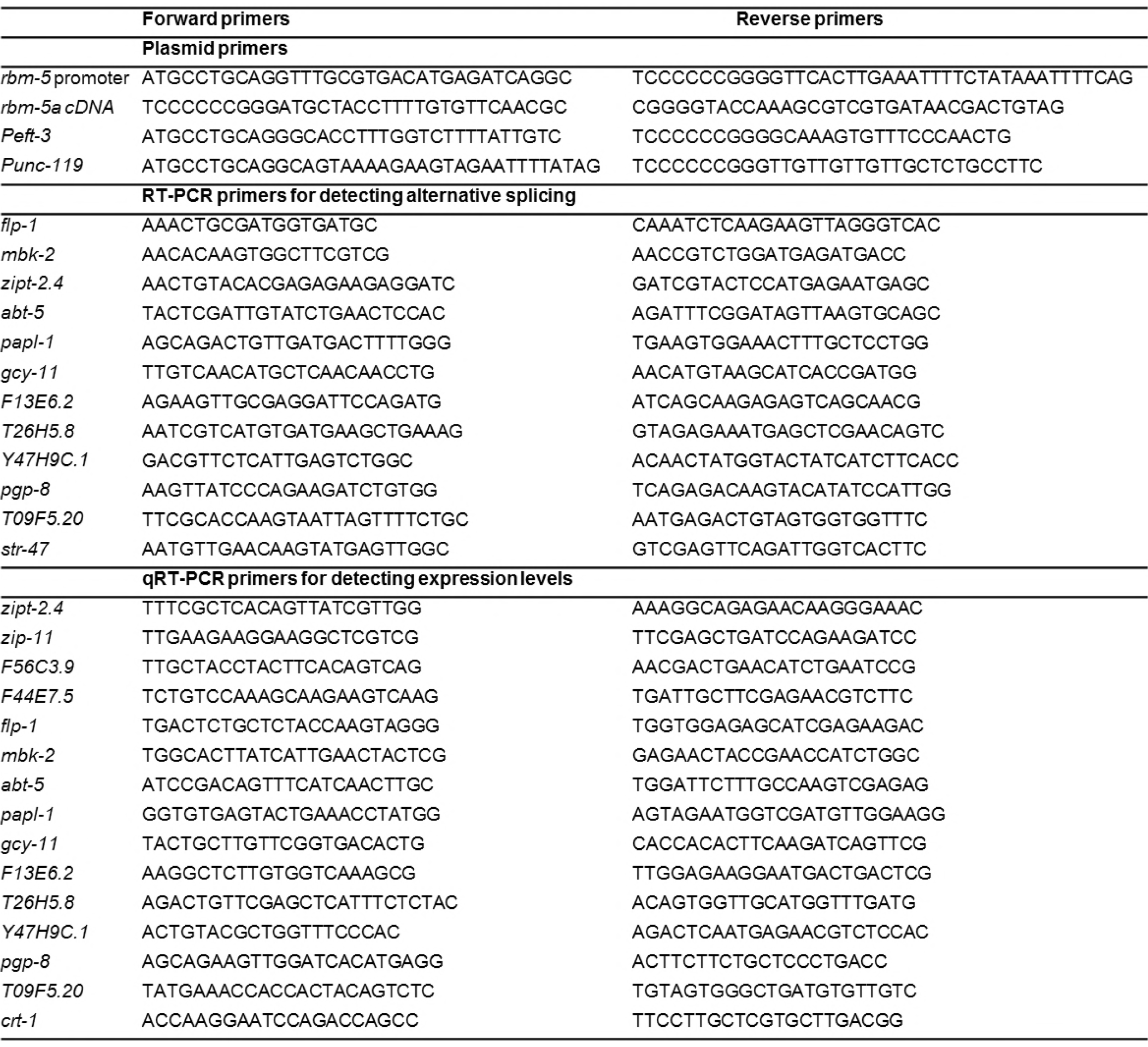
List of PCR primers.

Result S1: List of differentially expressed genes in *uaf-1(n4588)* and *rbm-5; uaf-1(n4588)* mutants.

